# Interspecies differences in proteome turnover kinetics are correlated with lifespans and energetic demands

**DOI:** 10.1101/2020.04.25.061150

**Authors:** Kyle Swovick, Denis Firsanov, Kevin A. Welle, Jennifer R. Hryhorenko, John P. Wise, Craig George, Todd L. Sformo, Andrei Seluanov, Vera Gorbunova, Sina Ghaemmaghami

## Abstract

Cells continually degrade and replace damaged and old proteins. However, the high energetic demand of protein turnover generates reactive oxygen species (ROS) that compromise the long-term health of the proteome. Thus, the relationship between aging, protein turnover and energetic demand remains unclear. Here, we used a proteomic approach to measure rates of protein turnover within primary fibroblasts isolated from a number of species with diverse lifespans including the longest-lives rodent, the naked mole rat and the longest-lived mammal, the bowhead whale. We show that organismal lifespan is negatively correlated with turnover rates of highly abundant proteins. In comparison to mice, cells from long-lived naked mole rats have slower rates of protein turnover, lower levels of ATP production and reduced ROS levels. Despite having slower rates of protein turnover, naked mole rat cells tolerate protein misfolding stress more effectively than mouse cells. We suggest that in lieu of rapid constitutive turnover, long-lived species may have evolved more energetically efficient mechanisms for selective detection and clearance of damaged proteins.

## Introduction

Protein homeostasis (proteostasis) encompasses an array of coordinated cellular functions that ensure the proper synthesis, folding and degradation of cellular proteins (1, 2). These functions are carried out by a number of cellular pathways that include the ribosomal translational machinery, the cellular chaperone network, the ubiquitin-proteasome system (UPS) and autophagic degradation (3). Combined, these pathways are required to maintain a functional proteome throughout the lifespan of an organism. A key feature of proteostasis is protein turnover, the process by which cellular proteins are continuously degraded and replaced by newly synthesized proteins (4). Because of its central role in protein quality control, there has been significant interest in understanding the relationship between alterations in protein turnover kinetics and proteostatic disruptions that occur during aging (1, 5–7).

Within a cell, proteins have widely varying half-lives ranging from minutes to years (8–10). Protein half-lives are established by a complex combination of intrinsic and extrinsic factors including degradative sequence motifs, the folding state of the protein, and activities of cellular degradative pathways (11–17). Protein half-lives are not only divergent within a proteome but also can vary for the same protein expressed in different organisms, tissues, cell types and environmental conditions (9, 18–21). It is thought that the constitutive clearance and replacement of cellular proteins accounts for the majority of their degradative flux within a cell (22, 23). In addition to this basal turnover, cells maintain a number of sophisticated pathways for specific detection and selective clearance of proteins that have been damaged through covalent modifications or detrimental conformational alterations (3, 24, 25). Together, the constitutive and selective clearance of proteins are important for maintaining a healthy proteome within a cell and alterations in protein turnover have been associated with the accumulation of damaged proteins and their pathogenic aggregation during aging and age-associated diseases (5, 6, 26–28).

Quantitative analyses of protein turnover have been significantly bolstered by recent technical advances in mass spectrometry-based proteomics. The use of stable isotope labeling in conjunction with LC-MS/MS has opened the door for analyses of protein turnover kinetics on a global scale (4, 9, 17, 18, 29). In a number of model systems, proteomic studies have shown that aged individuals have significantly slowed protein turnover compared to younger counterparts (30–32), although this effect may not be universal and appears variable for different proteins, tissues and organisms (33–36). More mechanistic studies have linked the age-dependent decrease in protein turnover and resulting loss in proteostasis to alterations in specific degradation pathways including the UPS and autophagy (35, 37, 38).

The above studies of aging organisms suggest that slower rates of protein turnover may be associated with less robust protein quality control and diminished proteostasis during aging. This trend is echoed in studies of invertebrates whose lifespans have been artificially prolonged by genetic alterations or life-extending treatments. For example, the ablation of the insulin-like growth factor 1 receptor *daf2* in *C. elegans* extends lifespan and results in an increase in global protein turnover rates, counteracting the natural decline in turnover that occurs during aging (31, 39–41). In apparent contrast, a number of studies have shown that protein turnover rates are actually reduced in mammalian model systems whose lifespans have been extended by dietary regiments, therapeutic interventions and genetic modifications (5). For example, mice that have been treated with rapamycin or calorie restriction, both well documented methods of lifespan extension, have reduced protein turnover rates (42–45). Similarly, in long-lived Snell dwarf mutant (*Pit1^dw^*) mice, rates of protein turnover are diminished (45). Thus, the nature of the relationship between protein turnover and longevity appears complex and remains enigmatic.

Within mammals, why are faster rates of protein turnover generally associated with youth, but organisms that are able to live longer have slower rates of protein turnover? To gain insight into this question, our laboratory previously conducted a comparative proteomic study of fibroblasts cultured from a number of rodent species with diverse lifespans and showed that within this limited set of organisms, longevity was correlated with slower rates of *in vitro* protein turnover (21). However, the generality and underlying basis of this trend were unclear. In the current study, we have expanded our cross-species comparisons of protein turnover kinetics to include mammalian species outside the order *Rodentia,* including the longest-lived mammal, the bowhead whale *(Balaena mysticetus)*. We show that the negative correlation between lifespan and protein turnover rates is apparent in mammalian species with lifespans ranging from 3 to 200 years. By comparing a representative pair of short-lived and long-lived species (mouse and naked mole rat, respectively), we investigated the ramifications of slower protein turnover on cellular ATP demand, generation of reactive oxygen species (ROS) and ability to survive protein misfolding stress. The results provide insights into the proteostatic costs and benefits of protein turnover and its relationship to longevity in mammals.

## Results

### Proteome-wide analysis of protein degradation rates (*kdeg*)

Skin fibroblasts were isolated from twelve different mammalian species and propagated in culture as previously described (Figure 1A) (21, 46). The collection included eight rodent species whose proteome-wide turnover kinetics were analyzed in a previous study (21). Additionally, the current study included four mammalian species: human (*H. sapiens*), cow (*B. bovine*), humpback whale (*M. novaeangliae*), and bowhead whale (*Balaena mysticetus*). The maximum lifespans of these organisms range from ∼4 years (mouse, hamster and rat) to ∼200 years (bowhead whale).

**Figure 1.**
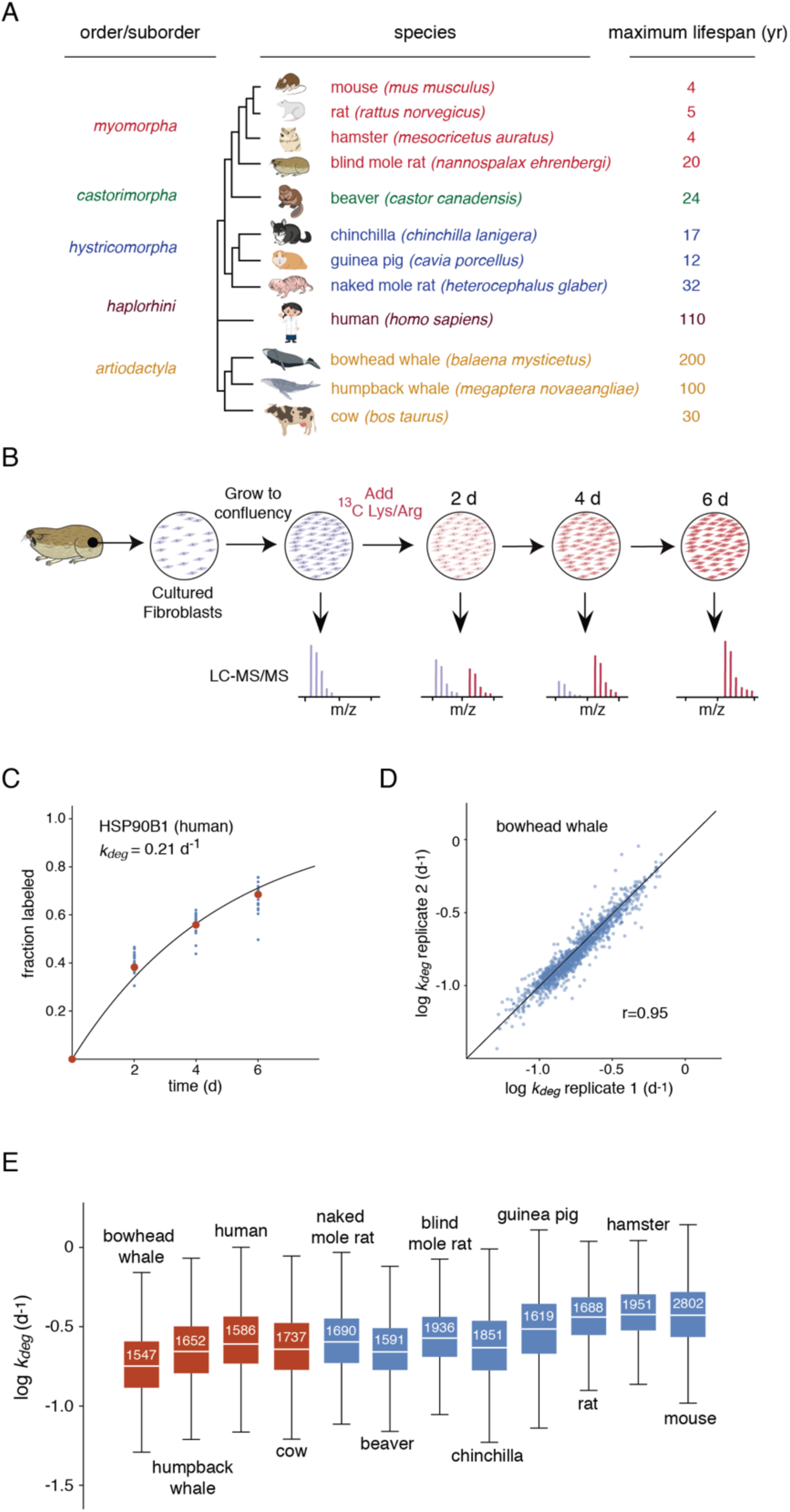
Proteome-wide quantitation of *k_deg_* in mammalian fibroblasts. A. Phylogenetic tree and maximal lifespans of species analyzed in this study. Colors indicate distinct orders and suborders. B. Dynamic SILAC experimental design. Blue and red colors indicate unlabeled and isotopically labeled spectra/cells, respectively. C. Labeling kinetics of human HSP90B1 (endoplasmin) shown as an example of protein-level determination of *k_deg_* values. Blue dots indicate fraction labeled for all peptides mapped to the protein and red dots indicated the median fraction labeled of all peptides. The blue curve is a least-squares fit to a first-order exponential equation with used to measure indicated *k_deg_* value. D. Pairwise comparison of *k_deg_* measurements in biological replicates of bowhead whale cells. The solid line indicates the identity line. The r value indicates the Pearson correlation coefficient. E. Distribution of *k_deg_* measurements for each species. The box plots indicate the median (white line), the interquartile range (box) and complete range (whiskers) of *k_deg_* measurements excluding far outliers (> 2 SD). The number in each box indicates the number of protein-level *k_deg_* measurements. Red and blue boxes distinguish data collected in this study and the previous study by Swovick et al. (21), respectively.

The global analysis of protein turnover rates was conducted using a dynamic (or pulsed) Stable Isotope Labeling in Cell Culture (SILAC) strategy (4, 9, 17, 18, 29, 47, 48) as illustrated in Figure 1B and described in detail in Materials and Methods. Briefly, cells were cultured at 37°C and grown to a contact-inhibited quiescent state prior to isotopic labeling. Typically, in dividing cells the fractional rate of isotopic labeling is influenced by both degradation rates and cellular proliferation rates (4, 49). By conducting the labeling experiments in a non-dividing state, we ensured that variations in cellular proliferation rates of different cultured cells were not influencing our observed labeling kinetics. Additionally, conducting the analyses in quiescent cells allowed the measurement of turnover kinetics for stable proteins whose half-lives are significantly longer than the doubling time of the cells (23, 49, 50).

The fractional labeling of tryptic peptides was measured after 2, 4 and 6 days of isotope incorporation with ^13^C-lysine and ^13^C-arginine. We were able to quantify the labeling for more than 6,000 peptides mapped to more than 1,500 proteins from each species (Table 1, Supplementary Table 1). For each protein, data from all corresponding peptides were aggregated and the median fractional labeling at each timepoint was calculated. In general, the labeling patterns of peptides mapped to the same protein closely mirrored one another (Supplementary Figure 1). For each protein, the median measurements of fractional labeling of mapped peptides at each time-point were fit to a single order exponential equation (assuming first order kinetics), and degradation rate constants (*kdeg*) were measured (Figure 1C, Supplementary Table 2). To assess the precision of the *k_deg_* measurements, we repeated the analysis for fibroblasts isolated from two individual bowhead whales (Figure 1D). The Pearson correlation coefficient for *k_deg_* measurements between these two biological replicates was 0.95. The distribution of *k_deg_* measurements for all identified proteins and for proteins shared between the species are shown in Figure 1E and Supplementary Figure 3A, respectively.

**Table 1.** Coverage of dynamic proteomic experiments. In order to compare an overlapping set of peptides shared between all species, all MS/MS data were searched against the mouse Uniprot database. “Unique peptides” indicates the number of unique peptide sequences. “Protein groups” indicates the number of homologous protein groups. The reported number of quantified protein groups is limited to those for which heavy to light (H/L) could be quantified by two separate spectra. The number of reported *k_deg_* values measured for protein groups is limited to those where two distinct peptides could be measured in two or more time-points and the least squares fit to a first-order equation had an R^2^ > 0.80. Numbers not in parentheses correspond to searches against species-specific sequence databases, or the most closely related sequence database available. Numbers in parentheses correspond to searches conducted against the *M. musculus* sequence database.

| Experiment | Time-point | Detected | | H/L quantified | | $k_{deg}$ measured | |
| --- | --- | --- | --- | --- | --- | --- | --- |
|  |  | Unique peptides | Protein groups | Unique peptides | Protein groups | Unique peptides | Protein groups |
| Human | 2d | 25881<br>(12370) | 4766<br>(3116) | 18523<br>(9585) | 2951<br>(1780) |  |  |
|  | 4d | 29539<br>(14008) | 5031<br>(3275) | 21149<br>(9934) | 2934<br>(1745) | 16091<br>(8018) | 2671<br>(1586) |
|  | 6d | 26394<br>(13418) | 4829<br>(3219) | 17458<br>(8889) | 2657<br>(1601) |  |  |
| Cow | 2d | 34025<br>(16372) | 4581<br>(3083) | 26322<br>(13019) | 3224<br>(2002) |  |  |
|  | 4d | 30458<br>(15243) | 4240<br>(2861) | 23136<br>(11656) | 2798<br>(1803) | 16978<br>(8578) | 2688<br>(1737) |
|  | 6d | 28758<br>(14387) | 4033<br>(2784) | 21605<br>(11042) | 2660<br>(1727) |  |  |
| Humpback Whale | 2d | 20932<br>(16245) | 3459<br>(3125) | 16014<br>(12559) | 2302<br>(1949) |  |  |
|  | 4d | 16932<br>(15165) | 3319<br>(2888) | 14735<br>(11438) | 2079<br>(1781) | 7926<br>(6181) | 1938<br>(1652) |
|  | 6d | 18584<br>(14507) | 3219<br>(2769) | 13687<br>(10722) | 1946<br>(1663) |  |  |
| Bowhead Whale rep 1 | 2d | 18136<br>(14027) | 3489<br>(2885) | 13733<br>(10686) | 2218<br>(1818) |  |  |
|  | 4d | 18324<br>(13882) | 3421<br>(2912) | 14022<br>(10675) | 2150<br>(1779) | 10194<br>(7795) | 1756<br>(1547) |
|  | 6d | 16781<br>(12877) | 3141<br>(2641) | 12734<br>(9775) | 2011<br>(1665) |  |  |
| Bowhead Whale rep 2 | 2d | 20237<br>(15371) | 3548<br>(3012) | 15532<br>(11898) | 2349<br>(1954) |  |  |
|  | 4d | 18628<br>(14347) | 3338<br>(1796) | 14426<br>(11070) | 2157<br>(1794) | 9866<br>(7675) | 1742<br>(1608) |
|  | 6d | 16336<br>(12744) | 3154<br>(2755) | 12281<br>(9668) | 1981<br>(1642) |  |  |

### Negative correlation between lifespan and protein degradation rates

Comparisons of *k_deg_* measurements between different pairs of species indicated relatively strong correlations with Spearman rank correlation coefficients (r_S_) values of all comparisons ranging from 0.63 to 0.87 (Supplementary Figure 2A). As expected, *k_deg_* values of closely related species were more correlated than evolutionary distant species (Supplementary Figure 2B). For example, the correlation of *k_deg_* measurements between bowhead whale and cow (55 MYA divergence, Figure 2A) was greater than that of bowhead whale and mouse, or bowhead whale and naked mole rat (87.5 MYA divergence, Figures 2B,C).

**Figure 2.**
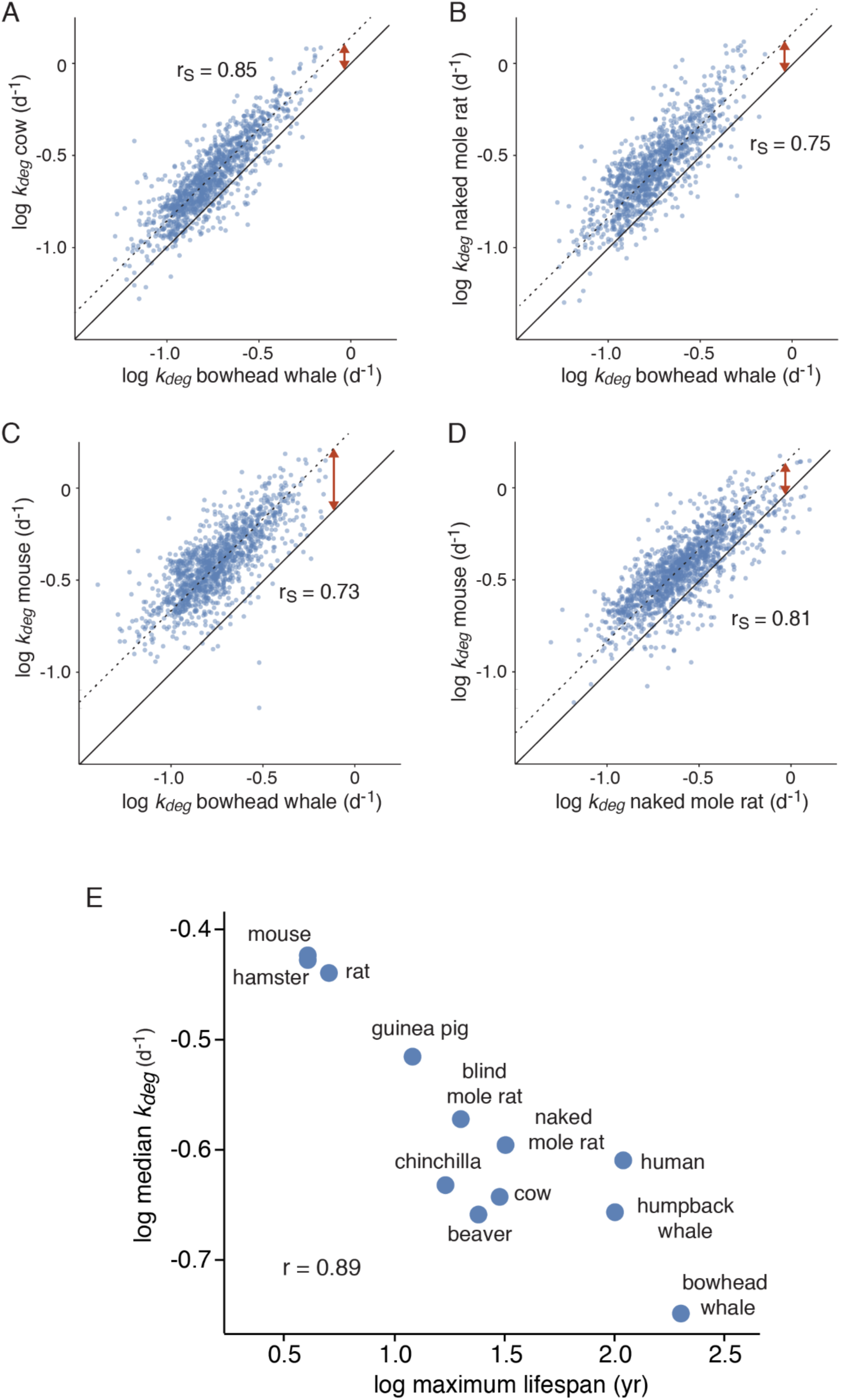
Cross-species comparison of *k_deg_* measurements. A-D. Pairwise comparisons of *k_deg_* measurements for representative pairs of species with variable lifespans and divergence times. Solid lines indicate lines of identity and dotted lines indicate lines of best fit. Red arrows highlight global shifts in distributions of *k_deg_*. r_S_ values indicate Spearman rank correlation coefficients. E. Correlation between species’ median *k_deg_* values and maximal lifespans. r value indicates Pearson correlation coefficient.

In addition to differences in correlation, pairwise comparisons of species highlighted global shifts in distribution of degradation rates (Figures 2A-D, red arrows). We calculated median degradation rates for proteomes of each species and assessed their correlation to several organismal properties including maximum lifespan and body mass. We observed a strong negative correlation between median *k_deg_* values and maximal lifespan taking into account all quantified proteins (Figure 2E, r=-0.89) or limiting the analysis to orthologous proteins shared between species (Supplementary Figure 3B, r=-0.90). A significant but weaker correlation was observed between *k_deg_* values and body mass (Supplementary Figure 3C, r=-0.76 and r=-0.78 for all and shared proteins, respectively). After correcting for the effect of phylogenetic distance (51), the correlation between median degradation rates and the maximal lifespans had a two-tailed p-value of 0.001, suggesting that these two parameters are independently correlated across a diverse set of mammals. Thus, our data indicate that longer lived organisms generally have slower global protein turnover rates.

### Degradation rates of abundant proteins are particularly diminished in long-lived organisms

We next assessed the relationship between organismal lifespan and degradation rates of individual proteins. We observed that this relationship was variable between proteins. For example, *k_deg_* measurements of orthologues of the kinase Ptk7 were relatively consistent among all analyzed species whereas *k_deg_* measurements of orthologues of the ribosomal protein Rps26 were significantly reduced in longer-lived organisms (Figure 3A). As a quantitative metric of lifespan dependence, we measured the slope of log *k_deg_* versus log lifespan for individual proteins (Figure 3A, Supplementary Table 3). We refer to this parameter as the turnover-lifespan slope (TLS). TLS values were calculated for all proteins where *k_deg_* measurements were determined for five or more species. We next determined if proteins with high or low TLS values were enriched in specific gene ontologies (GO) (Figure 3B, Supplementary Table 4). Among GO terms that were most prevalent within proteins with low TLS values (i.e. had more negative slopes) were cytosolic complexes such as the ribosome, proteasome and CCT/TRiC chaperonin, as well as proteins localized to the endoplasmic reticulum (ER) (Supplementary Figure 4A,B). Conversely, membrane proteins, mitochondrial proteins and proteins localized to intracellular vesicles such as the endosome had higher TLS values that were closer to zero.

**Figure 3.**
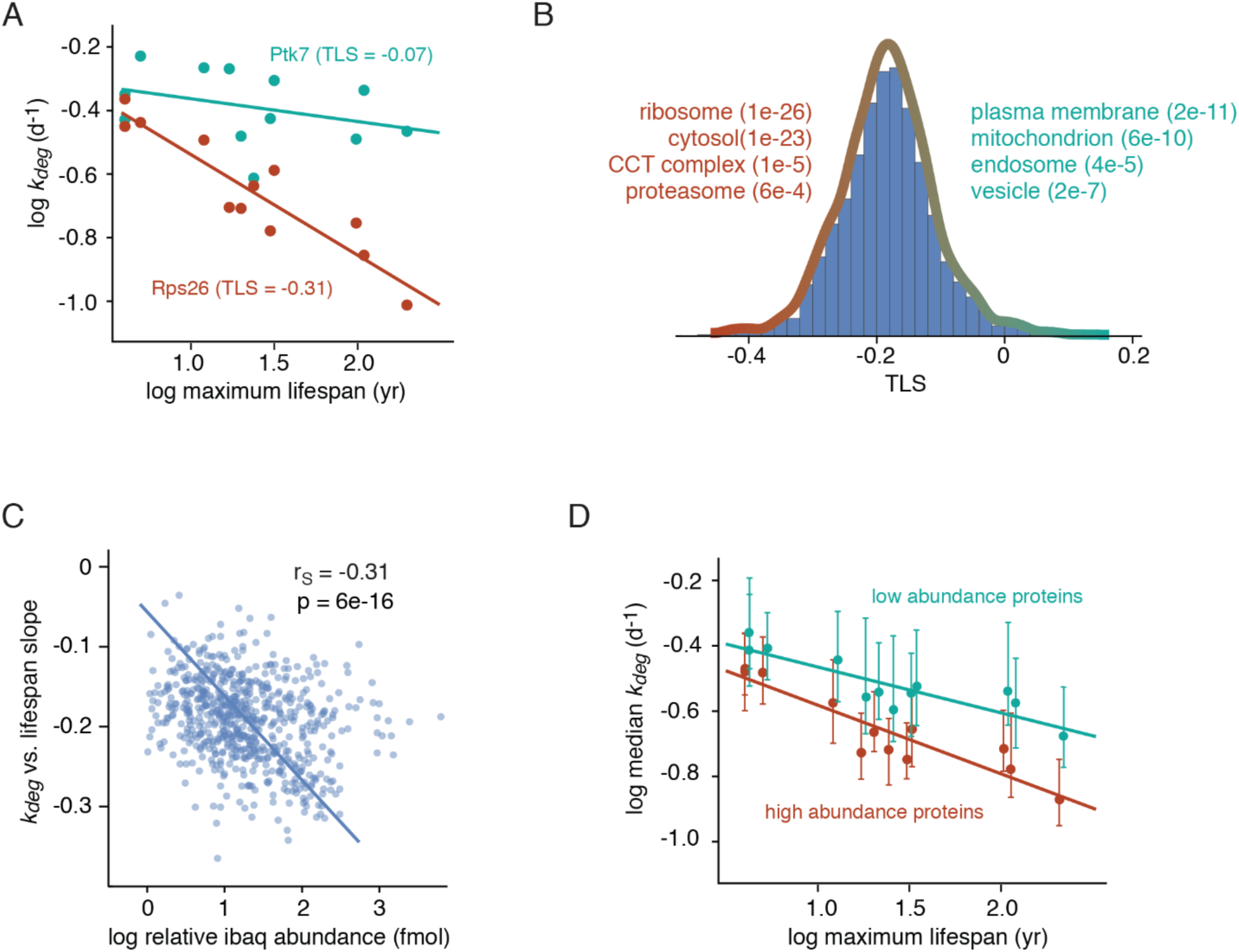
Differences in protein *kdeg* values between species are correlated with protein abundance. A. Correlation between *k_deg_* values of two example proteins (Ptk7 and Rps26) and maximum lifespans across species. The turnover-lifespan slope (TLS) measurements refer to slopes of the log-log plots. B. Distribution of protein TLS values. Steep and shallow TLS values are most enriched in proteins mapped to red and cyan GO terms, respectively. C. Correlation between protein TLS values and abundances (measured in mouse cells). Line indicates the line of best fit. r_S_ and p-values indicate the Spearman rank correlation coefficient and significance respectively. D. Correlation between median *k_deg_* values and maximum lifespans across species for the 500 most (red) and least (cyan) abundant proteins in the dataset.

We noted that the protein complexes whose degradation rates were most reduced in long-lived organisms were among the most highly abundant cytosolic proteins in eukaryotic cells (52–54). In order to determine if there was a general relationship between protein abundance and TLS, we measured the abundance of each protein within mouse cells using intensity based absolute quantification (iBAQ, Supplementary Table 5) (55). This proteomic methodology provides a proxy for the steady-state level of each protein by measuring the sum of MS peak intensities of all peptides matching the protein normalized by the number of theoretically observable peptides. By spiking in a set of absolute internal standards, we showed that iBAQ measurements were linearly correlated with absolute protein amounts across four orders of magnitude (Supplementary Figure 4C). Using this approach, we confirmed that proteins mapped to GO terms with the lowest TLS values were significantly more abundant than proteins mapped to GO terms with higher TLS values (Supplementary Figure 4D). Indeed, for individual proteins, there is a significant overall correlation between abundance and TLS (Figure 3C). As a further illustration of this phenomenon, we showed that the TLS values of the 500 most abundant orthologous proteins within our dataset were significantly lower than the 500 least abundant proteins (Figure 3D). Together, the data indicate that long-lived organisms may have evolved to specifically reduce turnover rates of highly abundant proteins.

### In comparison to mouse cells, naked mole rat cells have lower levels of ATP production

Protein turnover is one of the most energetically expensive processes in eukaryotic cells (56). Thus, the observation that turnover rates of abundant proteins are lower in long-lived organisms suggested that the longevity benefits of slow turnover may be related to reduced energy expenditure. We therefore explored metabolic differences between cells derived from a representative pair of species with divergent lifespans and protein turnover rates. We focused these studies on comparisons of mouse and naked mole rat cells as these two rodent species differ widely in lifespans and protein turnover rates yet are closely related evolutionarily and have similar body masses. Comparative studies of mice and naked mole rats have been used extensively in the aging field to identify cellular and physiological characteristics associated with longevity (57, 58).

We measured ATP production rates in quiescent mouse and naked mole rat cells using the Seahorse ATP Production Rate assay (59). The experiments provided measurements of oxygen consumption (OCR) and media acidification (PER) in real time. Under quiescent conditions, we observed that mouse cells consume oxygen and acidify the media at a more rapid rate than naked mole rat cells (Figure 4A). The addition of oligomycin, an ATP synthase inhibitor, results in a decrease in OCR that can be used to calculate the rate of ATP production from oxidative phosphorylation. The addition of a mixture of antimycin A and rotenone, inhibitors of electron transport chain complexes I and III, respectively, results in an increase in PER and can be used to calculate the rate of ATP production from glycolysis (59). Using these treatments, we observed that naked mole rat cells produce ATP at a slower rate from both glycolysis and oxidative phosphorylation in comparison to mouse cells (Figure 4B).

**Figure 4.**
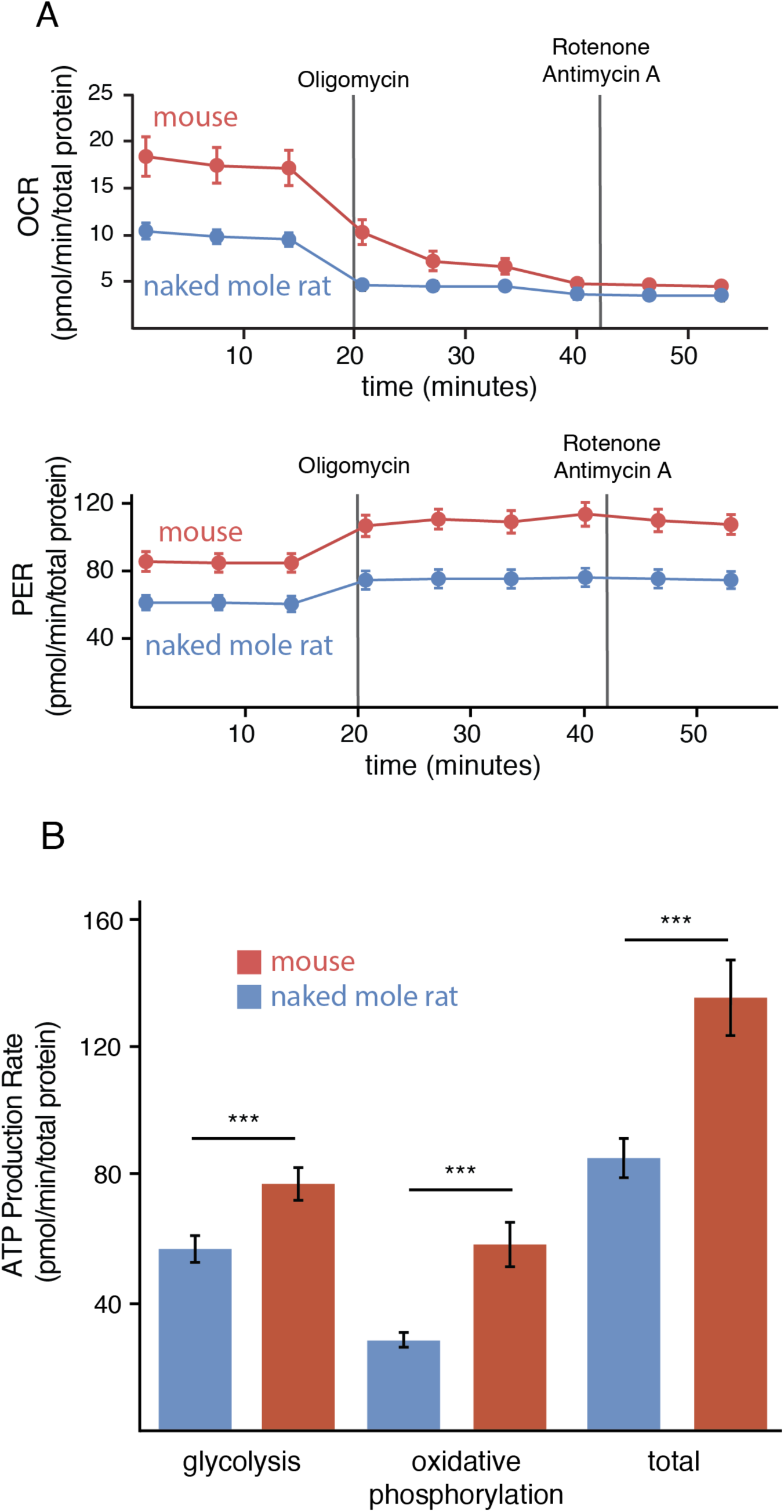
Differences in ATP production rates between mouse and naked mole rat cells. A. Rates of oxygen consumption (top) and proton efflux (bottom) of mouse (red) and naked mole rat (blue) fibroblasts. Solid vertical lines indicate the injection time of the specified inhibitor. B. Measurements of ATP production from either glycolysis or oxidative phosphorylation in mouse (red) and naked mole rat (blue) cells. Error bars indicate S.D. *** p-value < 0.0005.

In order to gain insight into potential molecular mechanisms of reduced ATP production rates in naked mole rat cells, we conducted label free quantitation (LFQ) proteomic experiments to identify proteins with altered steady-state expression levels relative to mouse cells. Quiescent mouse and naked mole rat cells were cultured in three biological replicate experiments, and relative changes in protein levels were analyzed using an LFQ methodology (See Materials and Methods, Figure 5, Supplementary Table 6). Proteins whose relative expression levels were lower in naked mole rats were most enriched in GO terms related to glycolysis and electron transport chain (ETC) proteins (Figure 5B, Supplementary Figure 5, Supplementary Table 7). Proteins with increased expression levels in naked mole rat cells had comparatively lower levels of enrichment for specific GO terms (Supplementary Table 7). The decreased expression levels of glycolytic and ETC proteins were confirmed by western blots (Figure 5C). The results suggest that lower rates of ATP production in naked mole rat cells may be related to lower expression levels of enzymes that play a role in the two major ATP producing metabolic pathways in eukaryotic cells (glycolysis and oxidative phosphorylation). These reductions are consistent with reduced total mitochondrial volumes within naked mole rat cells as determined by mitochondrial staining (Figure 5D).

**Figure 5.**
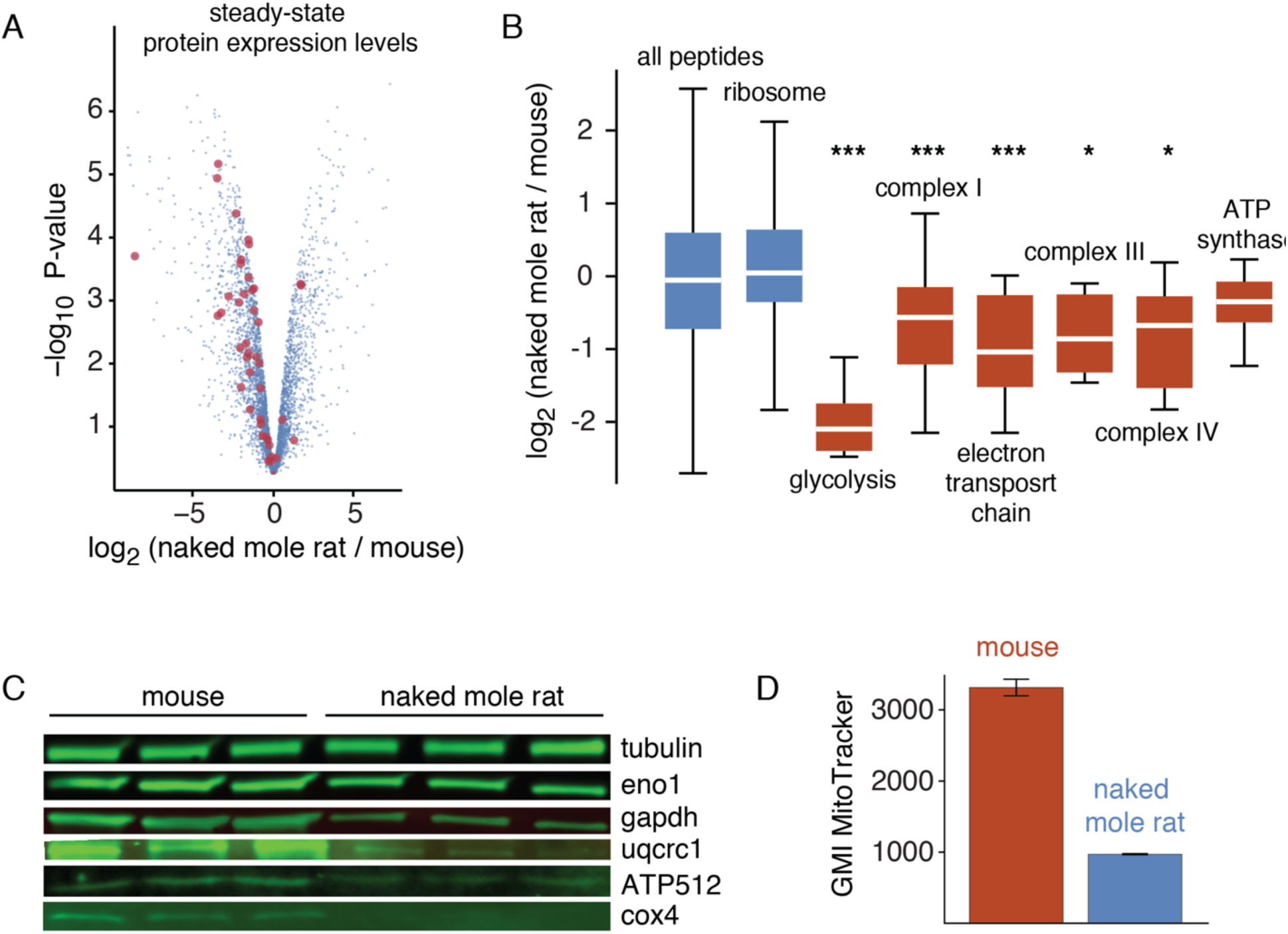
Proteome-wide differences in steady-state protein levels between mouse and naked mole rat cells. A. Volcano plot of p-value versus log_2_ ratio of expression levels in naked mole rat and mouse cells. Blue points represent individual proteins and red points highlight proteins involved in glycolysis and oxidative phosphorylation. B. Distribution of log_2_ expression level ratios for specified protein subsets. Box plot representations are as described in Figure 1. Red boxes highlight GO terms involved in ATP production. *, ** and *** indicate p-values of less than 0.05, 0.005 and 0.0005 in comparison to the global distribution using the Mann Whitney U test. C. Western blots of selected proteins from respiratory chain complexes (Uqcrc1, Atp512, Cox4) and glycolysis (Eno1, Gapdh). D. Measurements of geometric mean intensities of Mito Tracker mitochondrial staining of naked mole rat and mouse cells.

### In comparison to mouse cells, naked mole rat cells have lower levels of ROS production

Production of reactive oxygen species (ROS) is a byproduct of oxidative phosphorylation, and detrimental ROS-induced damage of cellular macromolecules accumulates during the course of aging (60). Given that quiescent naked mole rat and mouse fibroblasts have different rates of protein turnover and ATP production, we assessed whether they also have concordant differences in steady-state ROS levels. We measured cytoplasmic ROS levels using CellROX Orange, a cell-permeable probe whose fluorescence emission is enhanced upon oxidation (61). Quantitation of fluorescence by flow cytometry indicated that under quiescent conditions, mouse cells have significantly higher levels of cytoplasmic ROS than naked mole rat cells (Figure 6). The results are consistent with the idea that reduced energy expenditure due to slow protein turnover may be associated with reduced ROS accumulation in long-lived organisms.

**Figure 6.**
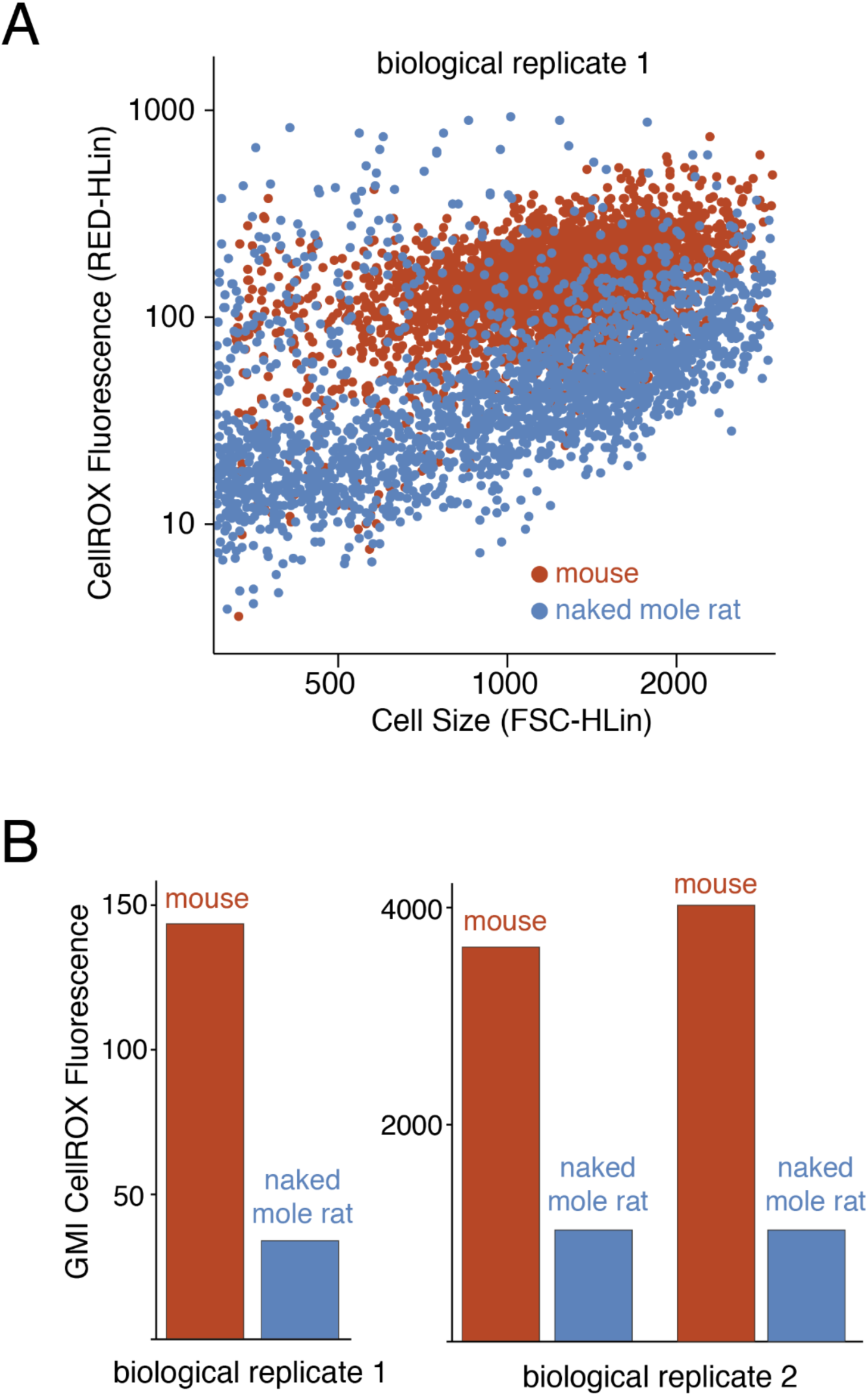
Differences in cellular ROS levels between mouse and naked mole rat cells. A. Flow cytometry analysis of CellROX fluorescence (ROS levels) and forward scatter (cell size) for mouse (red) and naked mole rat (blue) cells. B. Geometric mean intensities of CellROX fluorescence in mouse (red) and naked mole rat (blue) cells. Biological replicates represent cell lines cultured from individual organisms. The two pairs of measurements for biological replicates 2 represent measurements of distinct growths of the same cell line. The two biological replicates were measured using different flow cytometers as described in Methods.

### Slow protein turnover does not compromise proteostasis in naked mole rat cells

Slow protein turnover may be conducive to long lifespans because of concomitant reductions in energy expenditure and ROS accumulation. However, in theory, slow turnover may also have contradictory detrimental effects on proteostasis by reducing the clearance rates of damaged and misfolded proteins. We therefore assessed the ability of naked mole rat and mouse fibroblasts to tolerate proteotoxic stress. To induce proteotoxic stress, we treated cells with L-azetidine 2-carboxylic acid (AZC), a proline analog that can cause misfolding upon incorporation into proteins and is known to induce a cellular proteotoxic response (62–64). Mouse and naked mole rat cells were treated with varying concentrations of AZC in proline-containing media for 5 days and viability was assessed using Trypan Blue stain. Under dividing conditions, naked mole rat and mouse cells can tolerate similar doses of AZC before the onset of toxicity (Figure 7A). Interestingly, under quiescent conditions, naked mole rat cells can survive in the presence of higher concentrations of AZC in comparison to mouse cells (Figure 7B). Thus, slower protein turnover does not appear to comprise the ability of naked mole rat cells to respond to protein misfolding stress. In fact, under quiescent conditions, naked mole rat cells can tolerate higher concentrations of AZC in comparison to mouse cells.

**Figure 7.**
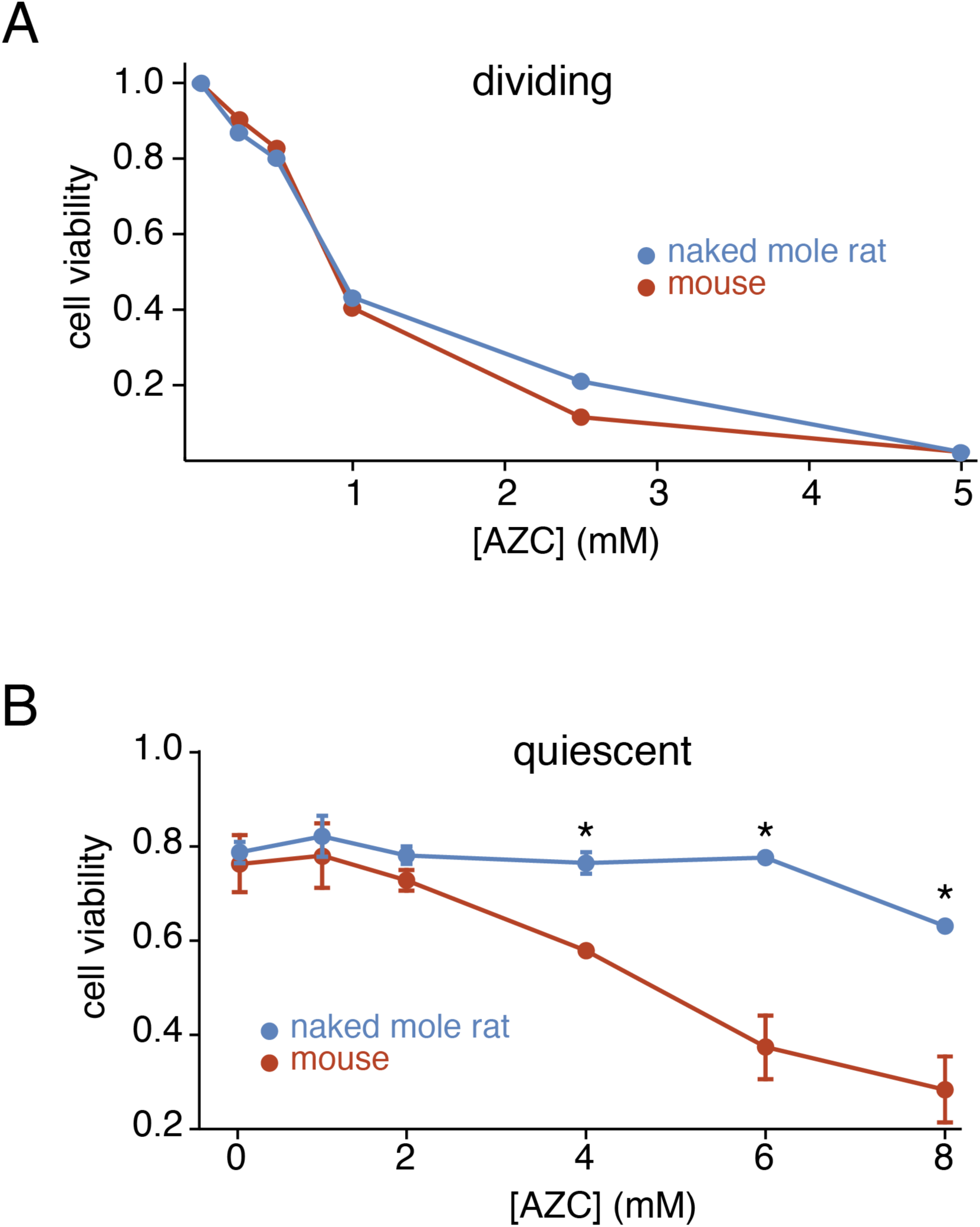
Cellular survival in presence of protein misfolding stress. A. Response to AZC treatment in dividing mouse (red) and naked mole rat (blue) cells. Cell viability represents the number of live cells compared to untreated. B. Response to AZC treatment in quiescent mouse (red) and naked mole rat (blue) cells. Cell viability represents the fraction of cells that excluded Trypan Blue. Error bars indicate S.D. * p-value < 0.05.

The enhanced ability of naked mole rat cells to tolerate AZC may either be due to lower levels of AZC incorporation into their proteomes or their ability to survive in the presence of higher levels of intracellular AZC-containing proteins. In order to distinguish between these two possibilities, we used a proteomic approach to quantify the levels of AZC-containing proteins in mouse and naked mole rat cells after exposure to AZC in the growth media (Figure 8A). Quiescent cells were treated with a non-lethal dose of AZC in proline-containing media and extracts were collected for LC-MS/MS analysis at varying exposure times. An initial pilot experiment indicated that AZC-containing peptides can be identified and quantified in AZC treated mouse cells with a low false positive identification rate (Figure 8B). The kinetic MS analyses quantified the level of AZC-containing peptides normalized with respect to their proline-containing counterparts. As a control, we also dosed cells with a benign isotopically labeled variant of proline (^13^C-proline) and quantified its fractional incorporation as a function of time. Together, the experiments allowed us to compare the accumulation of a benign and a proteotoxic analogue of proline in the proteomes of these two species.

**Figure 8.**
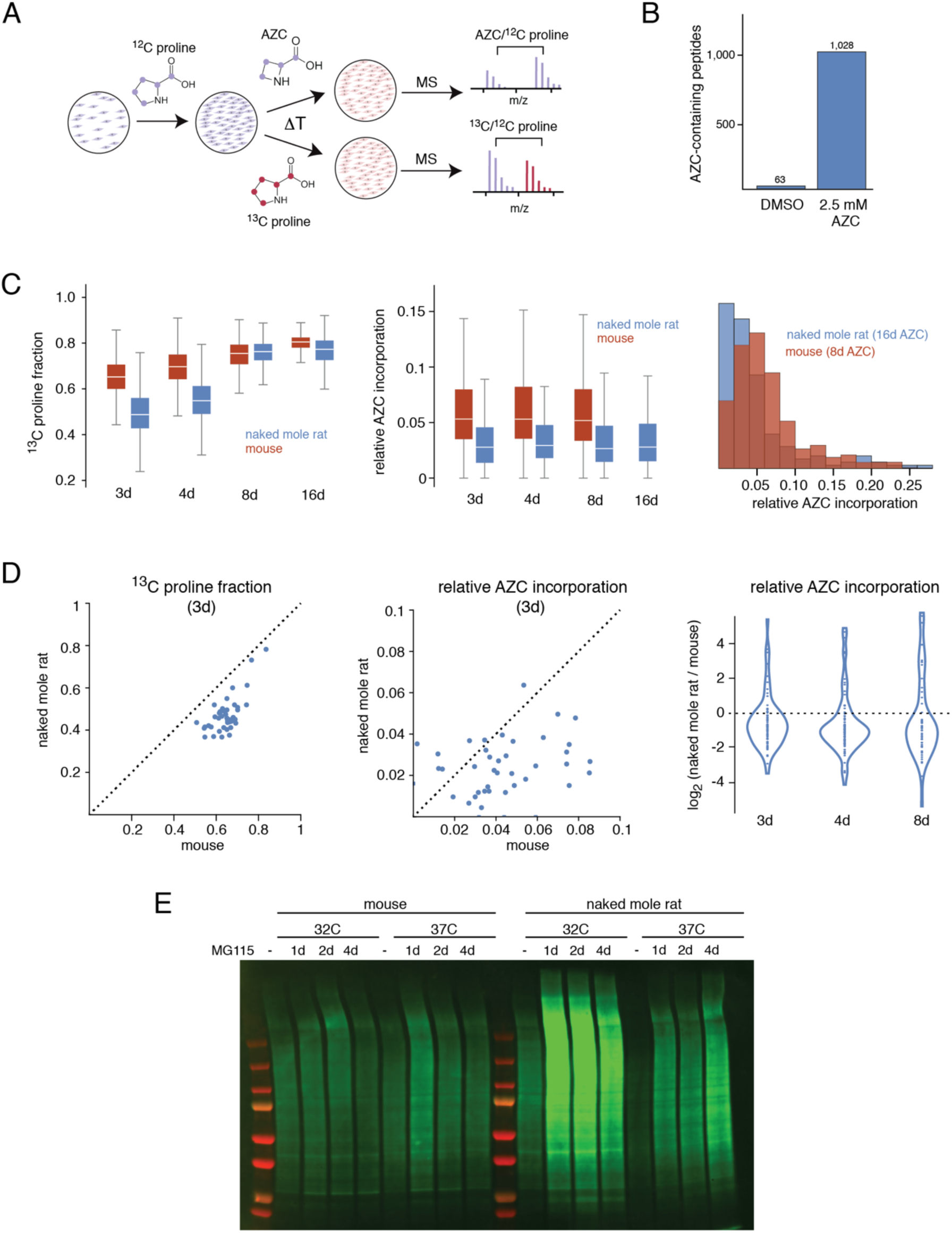
Accumulation of AZC in the proteomes of mouse and naked mole rat cells. A. Experimental design of proteomic experiments for measurement of AZC incorporation. Blue and red colors indicate ^12^C proline and either AZC or ^13^C proline, respectively. B. Number of AZC-containing peptides that were quantified in mouse cells treated with either DMSO or 2.5 mM AZC. C. Incorporation of ^13^C proline (left) and AZC (center, right) in mouse (red) and naked mole rat (blue) cells over time. Box plot representations are as described in Figure 1. The histogram to the right compares AZC incorporation in naked mole rats after 16 days of treatment with mouse cells after 8 days of treatment. D. Comparisons of ^13^C proline (left) and AZC (center, right) incorporation for individual peptides in mouse and naked mole rat cells. For pairwise comparisons (left, middle), peptides that could be quantified in both mouse and naked mole rat at 3d were compared. Similar plots for other timepoints are shown in Supplementary Figure 6. For the violin plot (right), log_2_ ratios of incorporation in mouse and naked mole rat cells are plotted for all peptides for which AZC incorporation could be measured in both species. E. Accumulation of ubiquitinated proteins upon proteasomal inhibition in mouse and naked mole rat cells. The experiment was conducted at 32°C and 37°C. Cells were treated with 5 µM MG115 proteasome inhibitor and extracts were collected at indicated times and analyzed by western blots using an anti-ubiquitin antibody.

The results indicated that mouse cells accumulated benign ^13^C-prolines in their proteomes faster than naked mole rat cells (Figure 8C, Supplementary Table 8). This observation was consistent with the faster rate of turnover observed in mouse cells using isotopically labeled lysines and arginines (Figure 2D). After 8 days of labeling with ^13^C-proline, the levels of incorporation were near 100% for both mouse and naked mole rat as their proteomes had almost fully turned over during that time. In contrast, levels of AZC incorporation remained significantly lower in naked mole rat cells even after 8 days of labeling. Indeed, naked mole rat cells exposed to AZC for 16 days contained less AZC in their proteomes than mouse cells exposed to AZC for 8 days (Figure 8C). Unlike naked mole rat cells, AZC-incorporation could not be measured for mouse cells at 16 days due to onset of toxicity and cell death at this time. The results suggest that the ability of naked mole rat cells to tolerate higher levels of AZC in the media is related to reduced accumulation of AZC-containing proteins in their proteomes.

We reasoned that there may be two possible explanations for why AZC-exposed naked mole rat cells accumulate less AZC in their proteomes in comparison to mouse cells. First, the ability of AZC to compete with proline during the process of protein synthesis (e.g. aminoacylation of tRNAs or incorporation into nascent polypeptides during translation) may be less efficient in naked mole rat cells. Second, naked mole rat cells may have an enhanced ability to specifically detect and clear AZC-containing and potentially damaged proteins once they have been synthesized. Our data appear more compatible with the second explanation as differences in levels of AZC-incorporation between mouse and naked mole rat cells are highly variable between orthologous proteins (Figure 8D). If naked mole rat cells had a reduced ability to aminoacylate tRNAs with AZC, or to utilize tRNAs charged with AZC during translation, we would expect a generally uniform reduction in AZC-incorporation for all observed proteins. Conversely, if misfolded AZC-containing proteins are cleared more efficiently in naked mole rat cells, we would expect variable levels of AZC-incorporation for different proteins (in comparison to mouse cells) as the effect of AZC-incorporation on misfolding would be dependent on its sequence context. In other words, AZCs incorporated in different proteins, or different parts of the same protein, are expected to have variable repercussions on folding, and hence different effects on clearance of the protein. When comparing AZC-incorporation for identical peptides between mouse and naked mole rat cells at different timepoints, we observe that the decrease in AZC-incorporation in naked mole rat cells is highly variable between peptides (Figure 8C, Supplementary Figure 6). These data suggest that the reduced accumulation of AZC-containing proteins in naked mole rat cells may be related to an enhanced ability to specifically detect and clear damaged proteins.

The best characterized mechanism for selective degradation of misfolded proteins in eukaryotic cells is ubiquitin-dependent proteasome degradation (65). To determine if the enhanced ability of naked mole rat cells to clear AZC-containing proteins was correlated to a higher activity of ubiquitin-dependent proteasomal degradation, we monitored the accumulation of ubiquitinated proteins following partial proteasomal inhibition (Figure 8E). The data indicate that the degradative flux of ubiquitinated proteins is higher in naked mole rats in comparison to mouse cells. More robust degradation of ubiquitinated proteins through the UPS may provide a mechanistic explanation for the ability of naked mole rat cells to selectively degrade damaged proteins.

## Discussion

To the best of our knowledge, the data presented in this paper represent the largest cross-species comparison of proteome turnover rates carried out to date. The *in vitro* analyses were conducted in quiescent dermal fibroblasts cultured from 12 mammalian species with widely divergent lifespans. The results highlighted a negative correlation between global protein turnover rates and organismal longevity. The generality of this correlation in other organisms, tissues, cell types, and environmental conditions remains to be determined. Nonetheless, the data provide a metabolic rationale for why it may be advantageous for longer lived organisms to maintain relatively slow rates of protein turnover.

Protein turnover is one of the most energetically demanding cellular processes in living organisms, accounting for as much as 25% of the total energy expenditure in some species (56). Thus, slower protein turnover rates can translate into diminished ATP demand and a concomitant reduction in the production of damaging reactive oxygen species (ROS) over the course of a long lifespan. A number of observations in this study are consistent with this idea. First, reductions in protein degradation rates in long-lived organisms are most pronounced for highly abundant proteins whose fractional turnover demands relatively more energy. Second, comparative analyses of mouse and naked mole rat cells (representatives of short-lived and long-lived species, respectively) demonstrate that the latter have slower rates of ATP production and lower expression levels of metabolic enzymes involved in glycolysis and oxidative phosphorylation. Third, naked mole rat cells have lower steady-state ROS levels in comparison to mouse cells. Although our analyses are necessarily correlative and do not directly demonstrate causation, the data are consistent with the idea that slower rates of protein turnover in longer lived organisms contribute to longevity by reducing energy demand and ROS production.

Our data also indicate that in naked mole rat cells, slow protein turnover does not diminish the tolerance for protein misfolding stress. Indeed, under quiescent conditions, naked mole rat cells have a higher tolerance for AZC, an amino acid analogue that results in misfolding upon incorporation into proteins. The data suggest that this tolerance may be related to the enhanced ability of naked mole rat cells to selectively degrade misfolded AZC-containing proteins. These observations are consistent with enhanced proteostasis in naked mole rat cells that have been reported in previous studies (66, 67).

How is it possible that naked mole rat cells have slow rates of basal protein turnover, yet have an enhanced ability to clear damaged proteins? It has long been appreciated that the turnover of the steady-state protein pool in a cell is largely stochastic and follows single-exponential first-order kinetics (23, 68). This trend has been largely confirmed by more recent proteome-wide analyses of turnover (4, 9, 17, 18, 29, 48, 55). From the perspective of total protein flux, it appears that for a given protein, any one molecule is as likely to be degraded than any other regardless of its age (elapsed time since its synthesis). The exception to this rule appears to be a fraction of newly synthesized proteins that are rapidly degraded immediately following translation and do not contribute to the steady-state protein pool in a cell (69). Thus, it appears that the vast majority of proteins that are present in a cell are not degraded because they have become damaged over time, but rather because their intrinsic properties have predetermined their basal half-lives. Nonetheless, protein damage is a common and dangerous occurrence in cells, and a number of sophisticated mechanisms have evolved for specific detection and removal of modified proteins (3, 24). In eukaryotic cells, these mechanisms include the ubiquitin proteasome system (UPS) and selective autophagy (28, 37, 65, 70). Thus, the total flux of proteins in a cell can be considered as having two components: nonselective basal turnover and selective clearance of damaged proteins. The former constitutes the larger component of total protein flux and is the rate that is typically measured by isotopic labeling experiments that quantify the overall fractional labeling of proteins over time.

Considering these two modes of protein flux, enhanced removal of damaged proteins can be achieved either by increasing the rate of nonselective basal turnover or deployment of more sophisticated surveillance systems for selective detection and clearance of aberrantly modified proteins. The former strategy is significantly more energy intensive and is likely to be associated with higher rates of ROS generation and associated damage over an extended lifespan. Whereas rapid nonselective basal turnover may represent a viable proteostatic strategy for short lived organisms, their associated costs in terms of ROS generation may not be conducive to longer lifespans. Thus, there may be selective evolutionary pressure for longer lived organisms to lower rates of basal turnover and, in lieu of reduced total protein flux, evolve more efficient mechanisms for selective clearance of damaged proteins.

The distinction between selective and nonselective protein clearance may also provide an explanation for the apparent contradiction in the relationship between turnover rates and longevity that has been observed in different model systems (5). In general, it appears that treatments and genetic modifications that extend the lives of invertebrates such as *C. elegans* and *D. melanogaster* model systems result in faster rates of total protein turnover (31, 39–41). Conversely, in mammalian models, life-extending treatments and genetic modifications are associated with slower protein turnover (42–45). If nonselective protein turnover plays a more prominent role in maintaining proteostasis in shorter lived organisms, it is understandable why extending lifespan may require faster rates of total protein flux. Conversely, in longer lived mammalian systems, increased ROS generation over time represents a greater impediment to longevity, and lifespan extension is associated with the lower energetic demands of slow protein turnover. In summary, our results provide support for the idea that the beneficial effects of protein turnover on proteostasis and longevity are mitigated by associated metabolic costs and liabilities.

## Methods and Materials

### Primary cell cultures

Dermal fibroblasts were isolated and cultured according to previously described protocols (21, 46). Briefly, the tissues were shaved and cleaned with 70% ethanol then washed with DMEM/F-12 medium (TheromFisher) with Liberase TM (Sigma) at 37°C on a stirrer for 15-90 minutes. Tissues were then washed and plated with DMEM/F-12 medium containing 15% fetal bovine serum (GIBCO) and Antibiotic Antimycin (GIBCO). When cells reach ∼80% confluency, isolated cells were frozen in liquid nitrogen within two passages. All subsequent cultures were grown in EMEM media supplemented with 15% fetal bovine serum (FBS), 100 U/ml penicillin, and 100 U/ml streptomycin at 37°C with 5% CO_2_ and 3% O_2_ except naked mole rat and whale cells which were cultured at 32°C with 5% CO_2_ and 3% O_2_. Prior to analyses, quiescent whale and naked mole rat cells were shifted to 37°C for 4 days.

### Isotopic labeling

Prior to isotopic labeling, cultures were grown to 100% confluency. Once cells ceased cell division and were contact inhibited, they were maintained in a quiescent state for 4 days. Subsequently, cells were acclimated to the labeling media (EMEM) supplemented with 15% dialyzed FBS (Thermo Scientific), 100U/ml penicillin, and 100 U/ml streptomycin for 4 days. After 4 additional days in adaptation media, the cultures were introduced to MEM media for SILAC (Thermo Scientific) supplemented with L-arginine:HCl (^13^C6, 99%) and L-lysine:2HCl (^13^C6, 99%;Cambridge Isotope Laboratories) at concentrations of 0.13 g/l and 0.0904 g/l respectively, 15% dialyzed FBS (Thermo Scientific), 100U/ml penicillin, and 100 U/ml streptomycin. After 0, 2, 4, and 6 days of labeling, cells were harvested, washed with PBS, and pellets were frozen before further analysis.

### Mass spectrometry sample preparation

Cells were lysed in 8 M Urea, 150 mM, and 50 mM HEPES (pH = 9.0). Cell pellets were resuspended in 50 µL of lysis buffer per 10^6^ cells and sonicated three times using a high-energy sonicator (QSonica, Newtown, CT) for 10s with 60s resting periods on ice. The samples were centrifuged for 5 min at 16000 x g and the supernatants were transferred to new Eppendorf tubes. Protein concentrations were measure using a bicinchoninic assay (BCA) kit (Thermo Scientific). Subsequent experiments were performed using 50 µg of total protein from each extract. Disulfide bonds were reduced with 5 mM Tris(2-carboxyethyl)phosphine (TCEP) Bond-breaker (Thermo Scientific) at RT for 1 hr, and proteins were alkylated with 10 mM iodoacetamide (IAA) at RT for 30 min in darkness. DTT was added to 1 mM to quench IAA and samples were diluted to a [Urea] of less than 1 M with 50 mM HEPES. To derive tryptic peptides, 1 µg of trypsin (selective cleavage on the C-terminal side of lysine and arginine residues) was added and the samples were incubated overnight at 37°C. To quench trypsin, formic acid was added to a final concentration of 1%.

To increase proteome coverage, high-pH fractionation was conducted on extracts before LC-MS/MS using homemade C18 spin columns. Eight different elution buffers were made with 100 mM ammonium formate (pH 10) (AF) with 5%, 7.5%, 10%, 12.5%, 15%, 17.5%, 20%, and 50% acetonitrile (ACN) added. After conditioning the column with ACN and AF, the samples were added and centrifuged. An AF wash was performed to remove any residual salt before the eight elutions were collected in fresh tubes. All fractions were then dried down and re-suspended in 10 µL of 0.1% TFA.

### Data analysis of isotopic labeling kinetics

MS2 data for all samples were searched against the appropriate Uniprot database using the integrated Andromeda search engine with MaxQuant software (71). For species where well-annotated sequence databases were not available, the data were searched against the closest related species. For example, data from humpback and bowhead whales were searched against the cow database. Additionally, for comparative analyses of shared peptide sequences, the data from all species were searched against the sequence database of *M. musculus* (22,305 entries, downloaded 8/7/2017). The maximum allowable numbers of missed cleavages in all searches were two and the FDR thresholds were set to maximum of 1%. SILAC peptide and protein quantifications were performed with MaxQuant using the parameter settings listed in Supplementary Table 9. For each peptide, heavy to light (H/L) SILAC ratio was determined by MaxQuant using a model fitted to all isotopic peaks within all scans that corresponding peptide spectral matches were detected. The H/L ratio for each peptide, obtained MaxQuant outputs, was subsequently converted to fraction labeled (H/H+L) measurements.

The determination of degradation rate constants (*k_deg_*) from the fraction labeled measurements were conducted in accordance to the kinetic model outlined previously (9, 17, 21). To obtain *k_deg_* measurements for each peptide, plots of fraction labeled as a function of time were fitted to a single exponential function using least squares fitting. To determine *k_deg_* at the protein level, the median fraction labeled measurements of all peptides mapped to specific proteins were fitted to a single exponential function using the least squares method. All reported *k_deg_* measurements at the protein level were required to pass three quality control criteria:

1. At least two unique peptide sequences were quantified in one or more time-points.
2. The fraction labeled at the protein-level was quantified in two or more time-points.
3. item The R^2^ for the fitted curve must be greater than 0.80.

Turnover-lifespan slopes (TLS) were measured as described in the Results section and gene ontology enrichment analyses were conducted on genes ranked based on their TLS values using GOrilla (73).

### Measuring steady-state protein levels

Three separate cultures were grown to quiescence in EMEM media supplemented with 15% fetal bovine serum (FBS), 100 U/ml penicillin, and 100 U/ml streptomycin. Cells were prepared for LC-MS/MS analysis as described above and LC-MS/MS analysis was performed as described below.

All samples were searched against the *M. musculus* Uniprot database using the integrated Andromeda search engine with MaxQuant software. Peptide and protein quantifications were performed with MaxQuant. All peptides that shared 100% sequence homology between mouse and naked mole rat were then used in subsequent analyses. Log_2_ fold change and p-values were calculated using the Proteus software (72) using the default settings and normalized by quantile normalization. Gene ontology enrichment analyses were conducted on genes ranked based on their Log_2_ fold change values using GOrilla (73).

To measure absolute protein levels, mouse fibroblasts were grown to quiescence at 37°C in EMEM media supplemented with 15% fetal bovine serum (FBS), 100 U/ml penicillin, and 100 U/ml streptomycin. The Universal Proteomic Standard (UPS2, Sigma-Aldrich) was dissolved in lysis buffer (8 M Urea, 150 mM, and 50 mM HEPES (pH = 9.0) and 4.24 µg was added to 13 µg of mouse lysate. The resulting mixture was prepared for LC-MS/MS analysis as described above and LC-MS/MS analysis was performed as described below.

MaxQuant protein intensities were divided by the number of theoretically observable peptides. The resulting iBAQ intensities of the standards were then log-transformed and plotted against the known log-transformed molar values of the standards. Linear regression was used to fit iBAQ intensities of the standards to the known molar protein amounts. From the resulting curve, the slope and y-intercept were used to convert iBAQ values to molar amounts for all identified proteins.

### Western blots

To perform Western blots, 20 µg of each extract was analyzed by electrophoresis in 10% polyacrylamide gels and transferred to polyvinylidene difluoride membranes using Trans-Blot SD Semi-Dry Electrophoretic Transfer Cell (Biorad). After 1 h incubation at room temperature in Odyssey Blocking Buffer (TBS) (Li-Cor Biosciences), the membrane was incubated with the indicated antibodies at 4°C overnight. The membranes were then washed with TBST/0.1% Tween 20, and the corresponding secondary antibodies were applied to the membranes for 1 h at room temperature. The membranes were then washed with TBST/0.1% Tween 20, and the detection of signal was performed using an Odyssey Scanning Instrument (Li-Cor Biosciences). The primary antibodies utilized and corresponding dilutions for the Western blots were Anti-alpha Tubulin antibody: 1:1000 (ab4074, Abcam), Recombinant Anti-ENO1 antibody [EPR19758]: 1:1000 (ab227978, Abcam), GAPDH Monoclonal Antibody (6C5): 1:1000 (AM4300, Invitrogen), UQCRC1 Monoclonal Antibody (16D10AD9AH5): 1:1000 (459140, Invitrogen), Recombinant Anti-ATP5L2 + ATP5L antibody [EPR15636]: 1:1000 (ab191417, Abcam), COX4 Monoclonal Antibody (K.473.4): 1:1000 (MA5-15078, Invitrogen) and Ubiquitin Monoclonal Antibody (P4D1): 1:1000 (3936S, Cell Signaling).

### ATP production rates

Two days prior to measuring ATP production rates, mouse and naked mole rat cells were plated onto a Seahorse XF96 Cell Culture Microplate (Agilent) at a cell density of 105,000 and 80,150 cells per well respectively to allow the cells to be 100% confluent. 24 hours prior to measuring ATP production, the media was changed to DMEM supplemented with 15% FBS, 100 U/ml penicillin, and 100 U/ml streptomycin. At the same time, an XFe96 Sensor Cartridge was hydrated with sterile H_2_O and placed in a non-CO_2_ 37°C incubator. On the day of measuring ATP production, 1 mL each of 1 M Glucose, 100 mM Pyruvate, and 200 mM L-Glutamine were added to XF DMEM. Cells were rinsed once with XF DMEM, cells were adapted to XF DMEM for 1hr prior to analysis. The sensor cartridge was incubated in a non-CO_2_ 37°C incubator for 1 hour with the sensors in XF Calibrant and 1.5 µM oligomycin added to port A and 0.5 µM Rotenone/Antimycin A added to port B. After 1 hour, the plate and sensor cartridge were loaded onto a Seahorse XFe analyzer according to the manufacturer and a XF Real-Time ATP Rate Assay was performed according to the company’s specifications. To normalize OCR and PER, the total protein levels in each well were calculated. To measure this, the total amount of protein was measured in triplicate for mouse and naked mole rat cells after culturing as described earlier. Cell number was counted for each culture, the mass of each culture was measured, and a BCA assay was performed to measure protein concentration. The resulting data was then used to calculate the protein concentration per cell. Assuming the majority of the cell is comprised of water (74), we calculated the volume of mouse and naked mole rat cells. Using the volume per cell and protein concentration per cell, we then calculated the total protein amount per cell. After completion of the ATP Rate Assay, the same plate was used to count the number of cells in each well. Cells were stained for 30 minutes with Hoechst 33342 (Cell Signaling Technologies) and imaged using a Celigo Imaging Cytometer (Nexcelcom). After imaging, the number of cells were counted in each well and using the protein amount per cell values we determined earlier, the total amount of protein present in each well was calculated.

### Quantifying cellular ROS

Mouse and naked mole rat cultures were grown to quiescence as described earlier and treated to a concentration of 5 µM of with CellROX Orange Reagent (Thermo Fisher) and incubated for 30 min at 37°C. After 30 minutes, the media was aspirated and cells were washed three times with warmed PBS. Cells were harvested using trypsin and centrifuged at 300 x g for 5 min. The resulting pellets were resuspended in 1 mL PBS and then transferred to a new tube diluted 1:2 with PBS. To quantify fluorescence, the Oxidative Stress program was run on a Muse Cell Analyzer (Luminex). To correct for auto-fluorescing of individual cells, fluorescence cells not treated with CellROX were analyzed. Thus, in cells treated with CellROX, only those cells whose fluorescence was higher than that of the background were measured. To ensure cellular debris and dead cells did not contribute to the overall fluorescence measurement, appropriate gates were used to exclude these cells from analysis. For analysis of biologically replicate cell lines (biological replicates 2 in Figure 6), confluent cells were treated with CellROX as described above, harvested and resuspended in PBS containing 1μg/ml DAPI (Thermo Fisher) for exclusion of dead cells and were kept in dark on ice until analysis. Flow cytometry analysis was performed on LSR II (BD). At least 20000 events were collected for each sample. FlowJo 7.6 (BD) software was used for data analysis.

### Mitochondrial staining

Mouse and naked mole rat cells were grown until confluency. Complete culture media containing 10 nM MitoTracker Deep Red (Thermo Fisher) was added to cells and cells were incubated for 30 minutes at 37°C. After incubation, cells were washed twice with PBS, harvested with trypsin and centrifuged at 300 x g for 5 min. Pellets were resuspended in PBS containing 1 μg/ml DAPI (Thermo Fisher) to exclude dead cells and immediately kept on ice before analysis. Flow cytometry analysis was performed on LSR II (BD). Dead cells and cellular debris were excluded from analysis by appropriate gating. At least 20000 events were collected for each sample. FlowJo 7.6 (BD) software was used for data analysis.

### AZC treatment and cell viability

To measure viability of dividing cells in response to AZC, cultures were grown in EMEM media supplemented with 15% fetal bovine serum (FBS), 100 U/ml penicillin, 100 U/ml streptomycin, and varying concentrations of AZC. Mouse cells were grown at 37°C while naked mole rat cells were grown at 32°C (their respective optimal growth temperatures). Two days after plating, cells were harvested and cell number was calculated. To measure cell viability, the total number of cells were compared to the number of cells counted in the plate treated no AZC.

To measure viability of quiescent cells in response to AZC, mouse and naked mole rat fibroblasts were cultured as described earlier and treated with varying concentrations of AZC (Sigma-Aldrich). After 5 days of treatment, cells were harvested and viability was determined by measuring Trypan Blue (Thermo Fisher) exclusion. To measure exclusion, 100 µL of a 0.4% trypan blue solution was added to 100 µL of cells and the resulting mixture was loaded onto a hemacytometer. To calculate cell viability, the number of blue cells was divided by the total number of cells and repeated four times and the average cell viability was then calculated for each sample.

### Measuring AZC incorporation

Eight total plates of both mouse and naked mole rat fibroblast cultures were grown to quiescence as described earlier. Four days after shifting quiescent naked mole rat cultures to 37°C, four plates of each species were treated with 2.5 mM L-proline (^13^C5, 99%; Cambridge Isotope Laboratories) and four with 2.5 mM AZC. After 3, 4, 8, and 16 days of labeling, cells were harvested, washed with PBS, and pellets were frozen prior to further analysis. Cells were prepared for and analyzed by LC-MS/MS as described above. All samples were searched against the *M. musculus* Uniprot database. To calculate the incorporation of ^13^C5 proline, the increase in fraction of labeled prolines were calculated as described above.

To measure AZC incorporation, a proline-specific modification with a mass change of −14.01565. was created within MaxQuant. To measure the relative level of AZC incorporation in each identified peptide, the following equation was used:

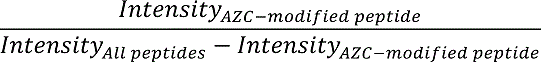

Where *Intensity_AZC-modified peptide_* is the intensity of identified peptides that have at least one AZC modification and *Intensity_All peptides_* is the intensity of all peptides quantified (with and without AZC) with the same sequence as the peptide in *Intensity_AZC-modified peptide_*. To compare global shifts in AZC levels over time, all peptides for which incorporation levels could be calculated in individual time-points were used. For pairwise-comparisons and Log_2_ ratio calculations, AZC incorporation of peptides with identical sequences were compared.

### Quantitation of ubiquitinated proteins

Mouse and naked mole rat cells were grown to quiescence at the appropriate conditions as described earlier (32°C for naked mole rat and 37°C for mouse). Upon reaching quiescence, cells were shifted to either 32°C or 37°C and allowed to acclimate for 4 days. Upon acclimation, cells were treated with 5 µM MG115 (Abcam). After 1, 2, and 4 days, cells were harvested and frozen prior to further analysis. Western blot analyses were conducted using an anti-ubiquitin antibody as described above.

### LC-MS/MS methods

To measure *k_deg_* of human, cow, humpback whale, and bowhead whale and label-free quantitation of mouse and naked mole rat, peptides were injected onto a homemade 30 cm C18 column with 1.8 um beads (Sepax), with an Easy nLC-1200 HPLC (Thermo Fisher), connected to an Orbitrap Fusion Lumos mass spectrometer (Thermo Fisher). Solvent A was 0.1% formic acid in water, while solvent B was 0.1% formic acid in 80% acetonitrile. Ions were introduced to the mass spectrometer using a Nanospray Flex source operating at 2 kV. Peptides were eluted off the column using a multi-step gradient that began at 3% B and held for 2 minutes, quickly ramped to 10% B over 5 minutes, increased to 38% B over 68 minutes, then ramped to 90% B in 3 minutes and was held there for an additional 3 minutes to wash the column. The gradient then returned to starting conditions in 2 minutes and the column was re-equilibrated for 7 minutes, for a total run time of 90 minutes. The flow rate was 300 nL/min throughout the run. The Fusion Lumos was operated in data-dependent mode with a cycle time of 3 seconds. The full scan was done over a range of 375-1400 m/z, with a resolution of 120,000 at m/z of 200, an AGC target of 4e5, and a maximum injection time of 50 ms. Peptides with a charge state between 2-5 were selected for fragmentation. Precursor ions were fragmented by collision-induced dissociation (CID) using a collision energy of 30 and an isolation width of 1.1 m/z. MS2 scans were collected in the ion trap with the scan rate set to rapid, a maximum injection time of 35 ms, and an AGC setting of 1e4. Dynamic exclusion was set to 20 seconds to allow the mass spectrometer to fragment lower abundant peptides.

Measuring incorporation of ^13^C proline and AZC in mouse and naked mole rat cells was done as above, with a few minor changes. The gradient was lengthened so that the total run time was 120 minutes, and the cycle time was reduced to 2 seconds. Dynamic exclusion was also increased to 45 seconds.

To perform iBAQ measurements in mouse cells, peptides were injected onto a similar column as above, with an Easy nLC-1000 HPLC (Thermo Fisher), connected to a Q Exactive Plus mass spectrometer (Thermo Fisher) using the Nanospray Flex source operating at 2 kV. Solvent A was 0.1% formic acid in water, while solvent B was 0.1% formic acid in acetonitrile using a flow rate of 300 nL/min. The gradient began at 3% B and held for 2 minutes, ramped to 8% B over 5 minutes, increased to 30% B over 68 minutes, ramped to 70% B in 3 minutes and held for 3 minutes before returning to starting conditions in two minutes. The column was re-equilibrated for 7 minutes, for a total run time of 90 minutes. The Q Exactive Plus was operated in data-dependent mode with a full MS scan followed by 20 MS/MS scans. The full scan was done over a range of 400-1400 m/z, with a resolution of 70,000 at m/z of 200, an AGC target of 1e6, and a maximum injection time of 50 ms. Once again, peptides with a charge state between 2-5 were selected for fragmentation. Precursor ions were fragmented by higher-energy collisional dissociation (HCD) using a collision energy of 27 and an isolation width of 1.5 m/z, with a 0.3 m/z isolation offset. MS2 were acquired with a resolution of 17,500 at 200 m/z, a maximum injection time of 55 ms, and an AGC setting of 5e4. Dynamic exclusion was set to 25 seconds.

## Associated Content

All raw and processed data are available in included Supplementary Tables (see below), and at ProteomeXchange Consortium via the PRIDE database (accession number PXD018325).

## Author Information

### Corresponding Author

*. Phone 585-275-4829

### Author Contributions

The study concept was conceived by K.S., V.G., A.S. and S.G. Its detailed planning was performed with contribution from all authors. Fibroblasts were isolated and cultured in the laboratory of V.G. and A.S. All wet-lab experiments were conducted by K.S. and D.F. Mass spectrometric analyses were conducted by K.S., K.W., and J.H. Data analysis was conducted by K.S. and S.G. The manuscript was written by K.S. and S.G. with input from all authors. All authors have given approval to the final version of the manuscript.

### Notes

The authors declare no competing financial interest.

## Acknowledgement

We thank the Alaska Eskimo Whaling Commission and Barrow Whaling Captains Association, for allowing us to sample bowheads; specimens were collected under NMFS Permit No. 21386 issued to the North Slope Borough Department of Wildlife Management, Utqiaġvik (Barrow), Alaska. We thank members of the Ghaemmaghami, Gorbunova and Seluanov labs at the University of Rochester for helpful discussions and suggestions.

## Supplementary Figures

**Figure S1.**
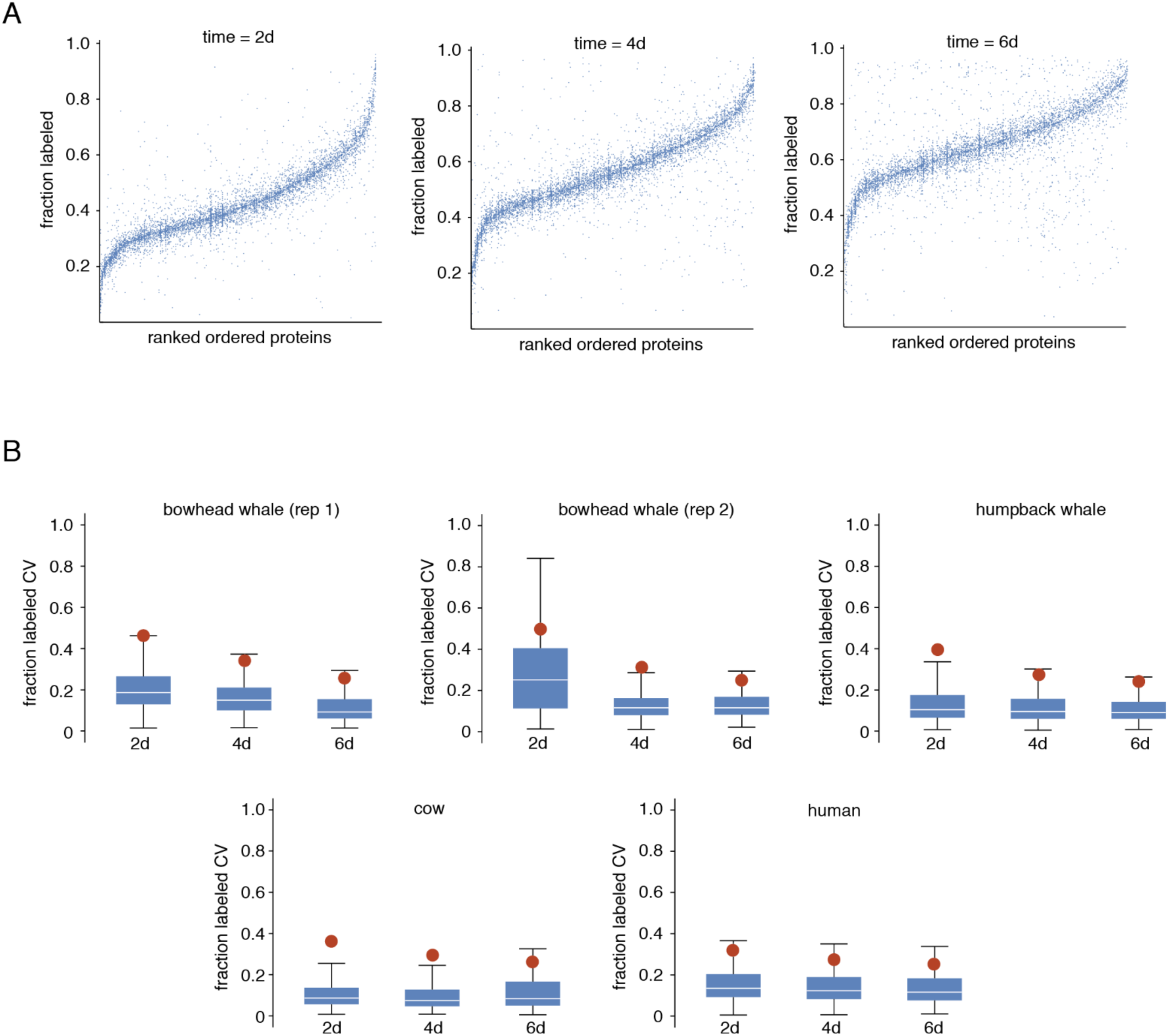
Peptide-level comparisons of fractional labeling in dynamic SILAC experiments. A. Rank-size distribution plots showing the fractional labeling of peptides matched to each protein within the proteome at different time-points. Vertical columns of blue points on the plot represent data for peptides matched to specific proteins. The plots generated for human data are shown as an example. Note that the range of measured fractional labeling for peptides within each protein is significantly narrower than the entire range of all measured peptides within the proteome. Plots from other species show similar trends. B. Box plots indicate the range of coefficient of variations (CVs) for measured fractional labeling of peptides mapped to the same protein. The dots indicate the CV for all peptides at a given time-point. Note that the intra-protein CV of peptide fractional labeling measurements are generally lower than the fractional labeling CV of all peptides in a dataset. Similar analyses for rodent species are presented in Swovick et al (21).

**Figure S2.**
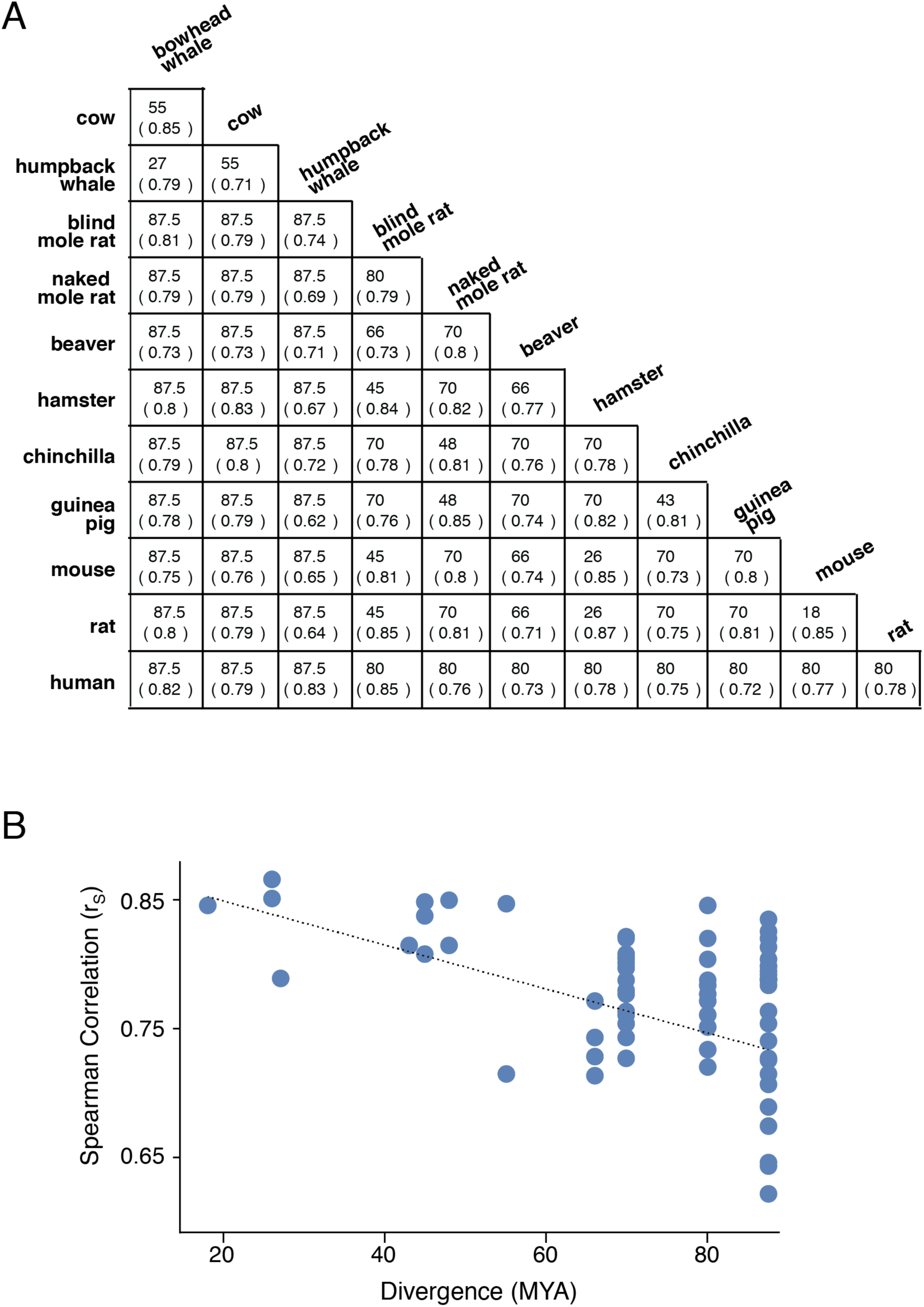
Cross-species correlations of protein *k_deg_* measurements decrease with increasing evolutionary divergence times. A. Divergence times in millions of years (top number) and spearman rank correlation coefficients of *k_deg_* values (bottom number) for pairs of species. B. Comparison of spearman rank correlation coefficients and divergence times for pairs of species. Dotted line represents the line of best fit.

**Figure S3.**
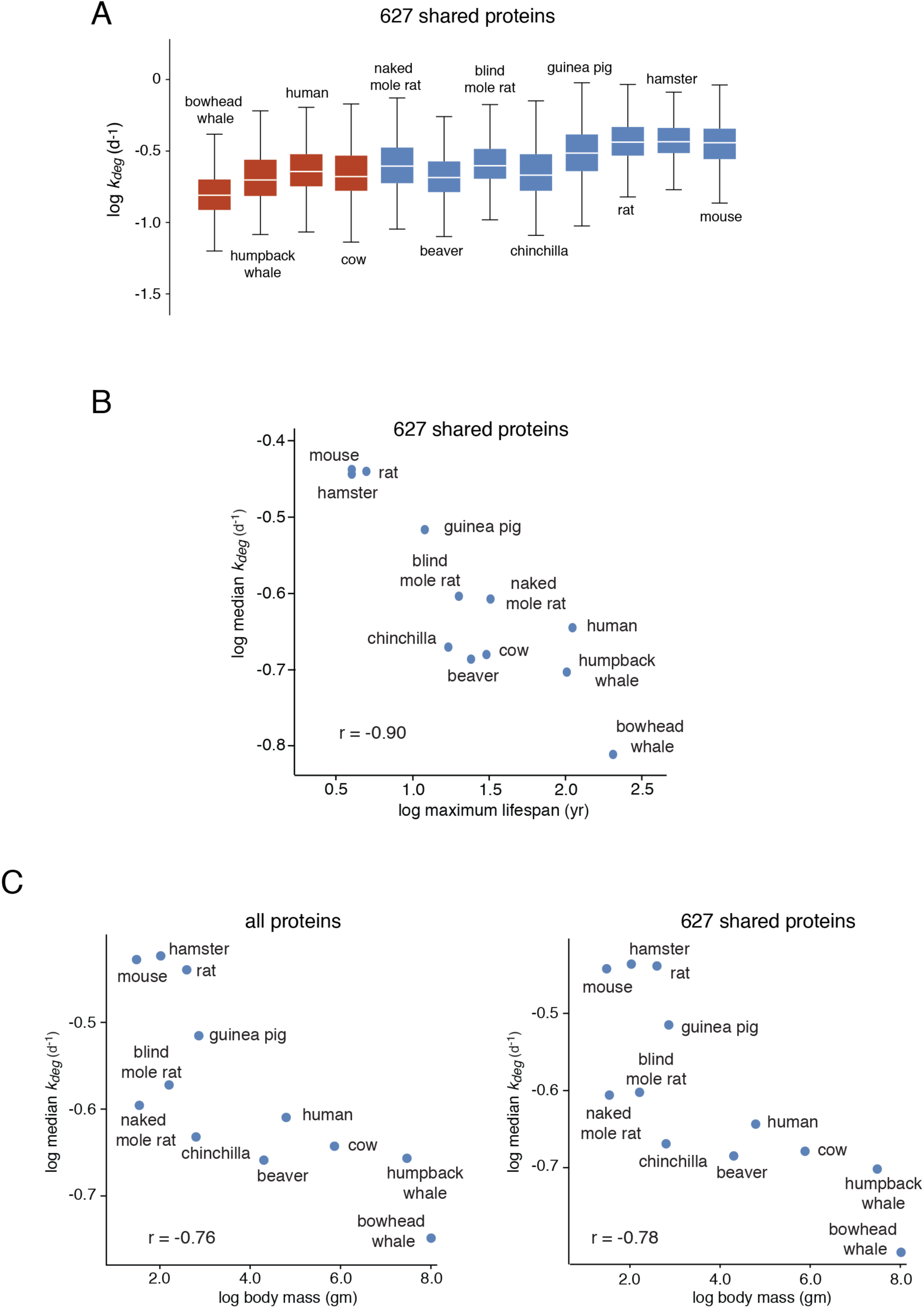
Cross-species comparison of *k_deg_* measurements for shared orthologues. A. Distribution of *k_deg_* values for shared 627 orthologous proteins quantified in all species. Red boxes indicate newly generated data while blue boxes indicate data generated by Swovick et al (21). B. Correlation of median *k_deg_* values with maximal lifespans for shared 627 orthologous proteins quantified in all species. C. Correlation of median *k_deg_* values with adult body mass for all measured proteins (left) and 627 orthologous proteins quantified in all species (right).

**Figure S4.**
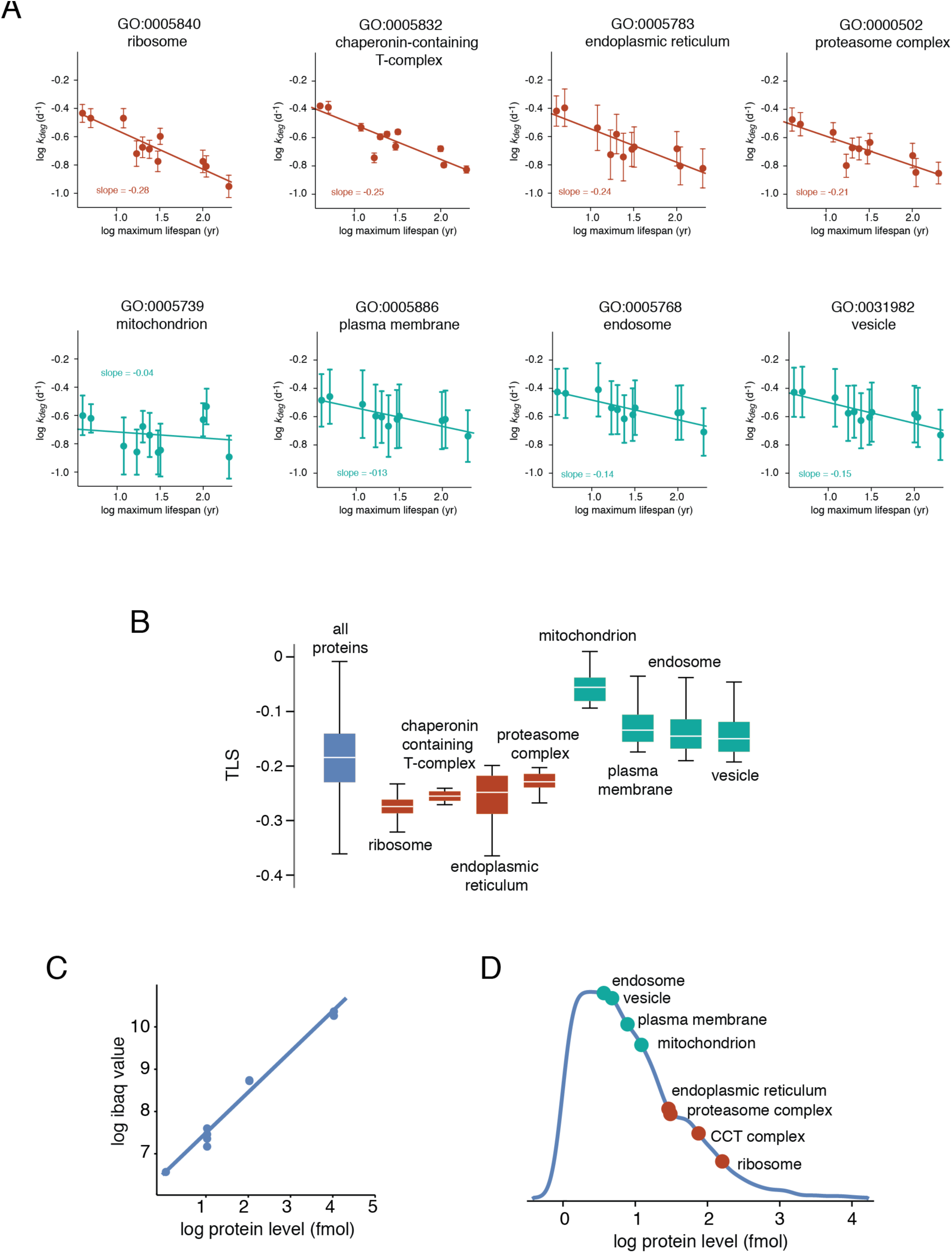
Cross-species differences in *k_deg_* are correlated with protein abundance. A. *k_deg_* of proteins mapped to specific GO terms in different species as a function of maximum lifespan. Red and cyan plots highlight GO terms that have steep and shallow slopes (TLS values), respectively. Error bars represent stand deviation of *k_deg_* measurements and lines represent lines of best fit. B. Distribution of TLS values for GO terms shown in A. Box plot representations are as described in Figure 1. C. iBAQ protein abundance calculations in mouse cells. (Left) Standard curve from spiked-in UPS2 standards in iBAQ experiment. (Right) Distribution of absolute protein levels in mouse cells. Indicated are the median abundances for GO terms listed in A and B.

**Figure S5.**
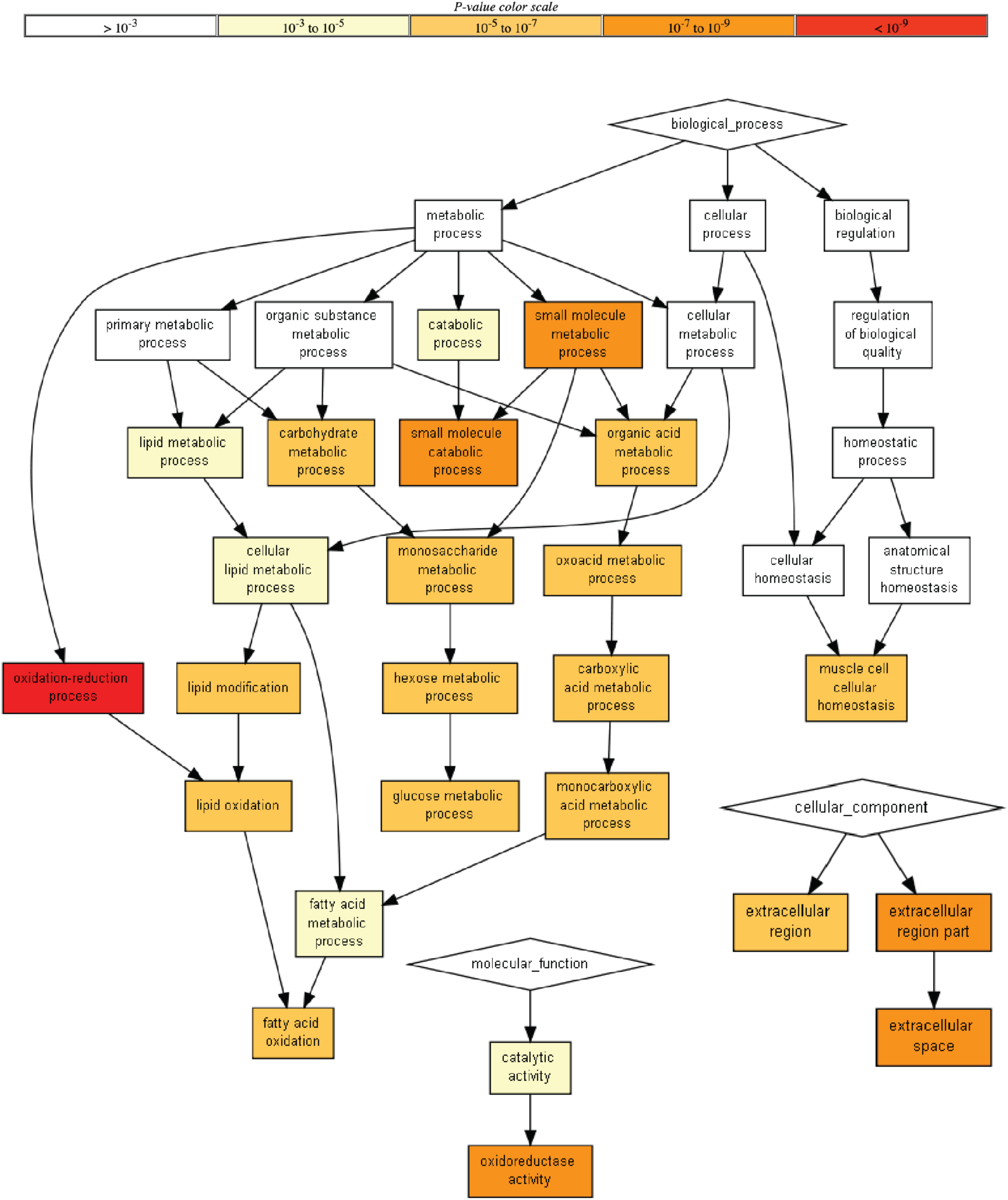
GO term enrichment analysis of protein level differences in mouse and naked mole rat cells. Network map of GO terms enriched within proteins that have higher expression levels in naked mole rat cells compared to mouse cells. The color scaling indicates p-values of the enrichment. The figure was generated using GOrilla (73).

**Figure S6.**
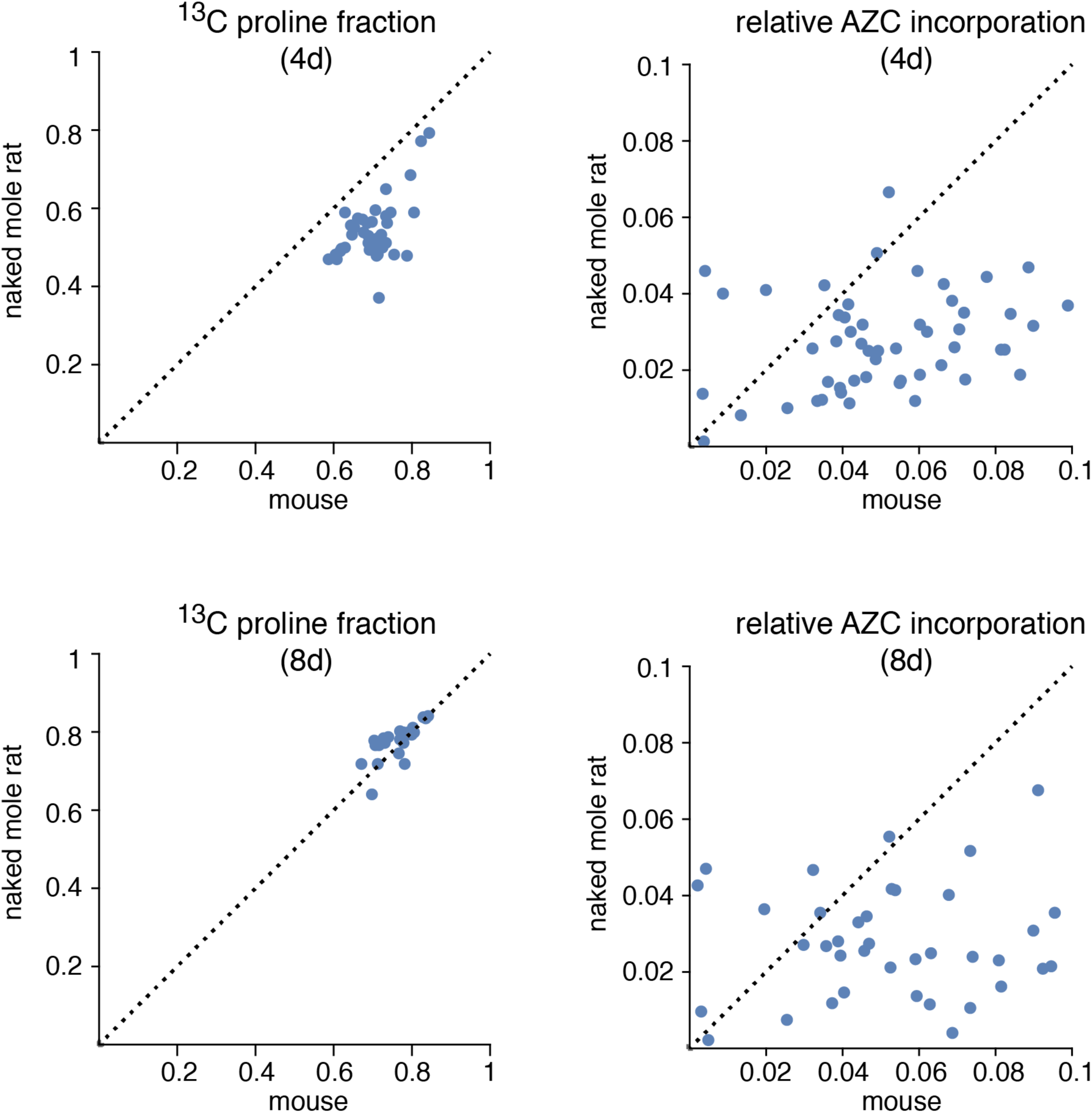
^13^C proline and AZC incorporation in mouse and naked mole rat cells. Pairwise comparisons of ^13^C proline (left) and AZC (right) incorporation for peptides quantified in both mouse and naked mole rat cells. Each point indicates a unique peptide quantified in both species on the specified day. Dotted line indicates the line of identity.

## Supplementary Tables

**Table S1. Dynamic SILAC LC/MS/MS data (TableS1.xlsx)**

**Table S2. *k_degradation_* measurements (TableS2.xlsx)**

**Table S3. Turnover-lifespan slope (TLS) measurements (TableS3.xlsx)**

**Table S4. Turnover-lifespan slope (TLS) GO enrichment (TableS4.xlsx)**

**Table S5. iBAQ abundance data (TableS5.xlsx)**

**Table S6. Mouse vs. naked mole rat LC-MS/MS label free quantitation (LFQ) data (TableS6.xlsx)**

**Table S7. Mouse vs. naked mole rat LC-MS/MS label free quantitation (LFQ) GO enrichment (TableS7.xlsx)**

**Table S8. AZC incorporation LC-MS/MS data (TableS8.xlsx)**

**Table S9 MaxQuant search parameters**

